# Taste preferences are influenced by temperature and the TRPM8 thermoreceptor

**DOI:** 10.64898/2026.02.18.706655

**Authors:** Kyle T. Zumpano, Chris Brooks, Christian H. Lemon

## Abstract

Oral temperature influences perceptual and behavioral responses to taste stimuli. Whether somatosensory processes contribute to this influence is unknown. Here we studied how stimulus temperature impacted mouse taste preferences and how these preferences were affected by the absence of TRPM8 (transient receptor potential melastatin 8). TRPM8 is a thermoreceptor supporting cool and warm temperature coding by somatosensory, including lingual trigeminal, neurons. Female and male TRPM8 gene deficient (KO) and C57BL/6J (B6) control mice (*n* = 69) entered thermolickometry tests that measured licking to two solutions of one taste chemical with different concentration-temperature properties across randomly interleaved brief-access (10-sec long) fluid exposure trials. Tastants were 15°C and 30°C bitter quinine (0.03 and 0.3 mM) or an appetitive sugar (sucrose or glucose, 100 and 500 mM). Mice were water-restricted or - replete during quinine and sugar tests, respectively, which were conducted separately and examined all possible concentration-temperature pairings across multiple days. Linear mixed model (LMM) analysis revealed that concentration and temperature both individually and additively affected licks emitted to quinine, with higher concentration and higher temperature (30°C) associated with the greatest reduction in quinine licks in B6 and KO mice. In contrast, LMM analysis indicated that temperature influenced licks to sugar only in KO mice, which made significantly fewer licks to higher than lower temperature sugars. Notably, KO mice also showed significantly reduced avidity to lick concentrated sugars than B6 mice. These data uncover novel roles for oral temperature and the TRPM8 thermoreceptor, and possibly its broader physiological influences, in taste preference behavior.

## 1. Introduction

Taste sensation, which supports the detection of nutrients and chemical toxins in ingesta, is modulated by diverse factors, including stimulus temperature. For example, rodent peripheral and central gustatory neural responses to oral presence of sucrose – an appetitive sugar taste stimulus – are enhanced by innocuous stimulus and oral warming and reduced by cooling (e.g., Breza et al., 2006; Lu et al., 2016; Talavera et al., 2005; Wilson & Lemon, 2014). Cold (10°C) fluid temperatures also suppress orosensory-guided licking of sucrose in water-replete rats compared to 20°C or 30°C (Torregrossa et al., 2012). Relatedly, human psychophysical data show the sweetness of sugars is enhanced by innocuous warming of taste solutions and lingual epithelia and suppressed by cooler temperatures (Bartoshuk et al., 1982; Calvino, 1986; Green & Frankmann, 1987; Nachtigal & Green, 2020), revealing an influence of temperature on gustatory perception.

Temperature effects on sugar taste responses are evinced to arise, in part, from thermal influences on taste receptor processes. Type II taste bud cells that transduce sweet-tasting chemicals utilize transient receptor potential (TRP) melastatin 4 (TRPM4) and 5 (TRPM5) ion channels downstream of ligand-receptor binding events to support depolarization and neurotransmitter release (Dutta Banik et al., 2018; Huang & Roper, 2010). TRPM4/5 is activated by increasing heat, with genetic knockout of TRPM5 impairing the ability of warming to facilitate gustatory nerve responses to sweet taste stimuli in mice (Talavera et al., 2005). Additionally, warming the tongue can elicit a thermal-induced sweet taste perception in humans (Cruz & Green, 2000) that is blocked by pharmacological inhibition of the sweet taste receptor TAS1R2/TAS1R3 (Nachtigal & Green, 2020), suggesting temperature change can directly affect taste receptor proteins.

While engaging with taste neurons, change in oral temperature causes substantially greater change in the excitability of thermosensory neurons of the trigeminal system (Lemon et al., 2016). Trigeminal neurons supply somatosensation to craniofacial and intraoral tissues and are insensitive to the oral presence of prototypical taste chemicals (Lemon et al., 2016; Simons et al., 2003). Primary trigeminal fibers express thermo-TRP channels implicated in thermosensory, and pain-related, transduction. These include the capsaicin- and heat-sensitive ion channel TRP vanilloid 1 (TRPV1) (Caterina et al., 1997) and the cool and menthol receptor TRP melastatin 8 (TRPM8) (Bautista et al., 2007; McKemy et al., 2002).

TRPM8 is excited (e.g., its current passage is increased) by temperature drops below about 30°C (McKemy et al., 2002; Tajino et al., 2011). Furthermore, TRPM8 is evidenced to influence warmth perception, as silencing TRPM8 impairs cutaneous detection of warm temperatures in mice (Paricio-Montesinos et al., 2020). This result associated with a loss of cooling-excited primary neurons that were inhibited by warmth in the genetic absence of TRPM8 signaling (Paricio-Montesinos et al., 2020). TRPM8 gene deficient mice also display deficits in responding to warmer oral temperatures, as they avoid licking 30°C and favor cooler water temperatures, like 15°C, in two-fluid choice tests whereas mice with intact TRPM8 function equally lick 30°C and 15°C water (Li et al., 2024). This behavioral difference associated with the appearance of anomalous positive correlation between trigeminal neural responses to oral presence of ∼30°C and aversively warmer temperatures in TRPM8 deficient mice, implying the valence of innocuous temperatures like 30°C can become negative without TRPM8 (Li et al., 2024).

While affecting the development of oral thermal preferences, TRPM8’s role in interactive temperature effects on taste is unstudied. Here, we used brief-access thermolickometry with TRPM8 deficient and control mice to test if fluid temperature and TRPM8 influence taste-guided licking of sugar (unconditionally appetitive) and bitter (innately aversive) taste stimuli. Our studies examined two fluid temperatures, 30° and 15°C, that show dependence on TRPM8 for oral sensation but are licked equally in water by wild-type mice (Li et al., 2024). We observed that when applied to aqueous solutions of bitter quinine, 30°C additively interacted with stimulus concentration to augment licking avoidance in both mouse lines. Moreover, 30°C suppressed licking towards sugars in TRPM8 deficient mice, which to our surprise also showed generally reduced avidity to lick sugar solutions. The latter result may reflect an adaptive response to a change in thermal perception and/or physiology associated with TRPM8 deficiency and implies thermosensation can influence taste preference behavior.

## 2. Methods

### 2.1. Mice

All experiments and procedures were approved by the University of Oklahoma Institutional Animal Care and Use Committee and followed the *Guide for the Care and Use of Laboratory Animals* (The National Academies Press). These studies used adult female and male C57BL/6J (B6; strain 000664, The Jackson Laboratory [JAX], Bar Harbor, ME) and homozygous TRPM8 deficient/knockout (KO; JAX strain 008198) mice. KO mice are derived from backcrosses with C57BL/6 mice (JAX). Sixty-nine mice entered behavioral tests. At the onset of behavioral training, mice were 17.7 (mean) ± 5.8 (SD) weeks of age. On average, female B6 mice (*n* = 17) weighed 19.7 ± 1.3 g and males (*n* = 18) 26.6 ± 2.0 g. Female KO mice (*n* = 17) weighed 20.8 ± 1.6 g and males (*n* = 17) 28.7 ± 2.3 g.

Before experiments, mice were group housed with *ad libitum* access to standard mouse chow and filtered water. Once the experiment started, each study mouse was singly housed with *ad libitum* food access, but water availability was regulated as below. Mouse cages were of the disposable type and were housed in a ventilated (HEPA filtered) rack system (Innovive, San Diego, CA) located in a climate-controlled (∼22°C ambient temperature) room maintained on a 12:12 hour light:dark cycle. Individual cages were prepared with standard bedding and enrichment materials (e.g., paper huts, cardboard tubes). Cage bedding was changed on a regular schedule prior to experiments, and only when needed during studies to mitigate housing disruptions.

### 2.2. Experimenter blinding

Prior to experiments, a unique alphanumeric code was assigned to each study mouse and used to label their cage in lieu of all other identifying information. Study mice cage locations in the colony housing rack were then randomized. These procedures were performed by lab personnel not directly involved with data collection, which blinded the experimenter handling and running mice to mouse strain (i.e., B6 or KO). All mice had black coats and indistinguishable outward appearances, facilitating blinding. Blinding was used to collect behavioral data across all training and test sessions. The experimenter was unblinded to the squad undergoing tests only after all data collection was complete.

### 2.3. Water restriction schedule

An overnight water restriction schedule was imposed on mice the day before their lickometer training commenced. This schedule aimed to motivate mice to lick fluids offered during fluid exposure training sessions. To do this, the water bottle for each home cage was removed and replaced with a marble-weighted bottle with no water. This weighted bottle blocked the water access hole on the cage top. Water restriction conditions were also used during tests with quinine. For sugars, water restriction was implemented during initial sugar orientation sessions but not during sugar tests, as below. The intent was that mice would consume their daily fluids in a lickometer when performing in training and test phases under water restriction. However, individual mice were given an additional ∼45-min access to water in their home cage following a brief-access session if their daily measured body weight fell below about 80% of their baseline weight.

### 2.4. Lickometer apparatus

Lickometer training sessions were partly carried out using standard “Davis Rig” contact lickometers (DiLog Instruments, Tallahassee, FL; Med Associates, St. Albans, VT). The Davis Rig can record licking responses made by a mouse to different fluids offered on sequential, seconds-long trials while randomizing the order of fluid presentation within each test session. The short trial length and limited number of fluid trials offered during brief-access tests aimed to mitigate post-ingestive influences (e.g., satiation) on licking responses and capture ingestive/licking behavior driven by oral sensation (Boughter et al., 2002; Smith, 2001).

Mouse licking responses to temperature-controlled fluids were measured using a custom-modified Davis Rig, as described (Li et al., 2024). Briefly, our “thermolickometer” functioned like a normal Davis Rig contact lickometer, capturing lick rates to different fluid sipper tubes randomly offered to mice on discrete trials, but held the temperature of each fluid at a specified set point value. During all temperature-controlled brief-access sessions, actual fluid temperatures were recorded before and after each test by placing a small thermocouple probe just inside the sipper tube tip.

### 2.5. Lickometer training

All brief-access tests were preceded by a training period, which oriented mice to the apparatus and brief-access procedure. Overnight water-restricted mice were individually trained to receive fluids in a standard Davis Rig over four days. On training days 1 and 2, mice were allowed free access to one sipper tube filled with room temperature purified water for 30 min to habituate them to receiving fluids in the rig chamber. The 30 min session started with the first lick on the sipper tube. Days 3 and 4 familiarized mice with the brief-access fluid exposure procedure by offering them different sipper tubes of room temperature purified water over 20, 10-sec access trials. Once a tube was presented, mice were allowed 30 sec to make a lick, which started the trial. Zero licks were recorded if no licks were made after 30 sec. Inter-trial intervals were 10 sec. Mice typically showed avid water licking during brief-access sessions on days 3 and 4; these data were also used as baseline responding for quinine tests, as below. Per-trial averages (mean water licks in 10 sec) were similar between lines: B6, 49.9 ± 1.4 SEM; KO, 46.1 ± 1.4 SEM (n.s. effect of mouse line, *F*_(1,64)_ = 3.9, *p* = 0.053; n.s. mouse line × sex interaction, *F*_(1,64)_ = 1.7, *p* = 0.203; two-way ANOVA). After completing the last day of training, mice were returned to their single-housing cage and given *ad libitum* access to water for approximately two days, prior to starting tests with taste stimuli.

### 2.6. Brief-access ffuid exposure tests

After this break, mice returned to an overnight water-restriction schedule and entered brief-access fluid exposure tests with temperature-controlled taste stimuli in the thermolickometer. Three taste stimuli were examined: the bitter stimulus quinine-HCl (quinine), the disaccharide sucrose, and the monosaccharide D-glucose (glucose). Instead of only one sugar, sucrose and glucose were tested given evidence of functional differences between them, including differences in signaling pathways (Ascencio Gutierrez et al., 2024; Breslin et al., 2021). All chemical stimuli were high purity (Sigma-Aldrich, St. Louis, MO) and dissolved in purified water.

Quinine, sucrose, and glucose were tested separately, on different weeks, and on different mice. Because all mice could not be tested at once, two separate squads of mice were each used for quinine, sucrose, and glucose tests. Across squads and stimuli, the general format of the daily test sessions was the same: two solutions of one taste chemical that differed in concentration (two levels) and/or temperature (two levels) were proffered over a series of short exposure trials where lick activity was recorded. There were four concentration-temperature configurations of each tastant, with all possible pairs of these configurations presented across six separate tests, as follows.

#### 2.6.1. Quinine experiment

Two squads of water-restricted (thirst-motivated) mice received the six different quinine concentration/fluid temperature test conditions across six sequential days (**Table 1**). Water restriction is needed to motivate mice to lick bitter quinine solutions (e.g., Hsiao & Fan, 1993). All mice within a squad were generally tested on the same days, with test order across days randomized for each mouse using a Latin square design. This counter balanced test order across subjects to control for a potential longitudinal effect of exposure, which was statistically evaluated as below.

**Table 1.** Stimulus concentrations and temperatures used in two-fluid brief-access thermolickometry tests with quinine.

| Test | Fluid 1: concentration, temperature | Fluid 2: concentration, temperature |
|---|---|---|
| 1 | 0.03 mM (low), 15°C (low) | 0.03 mM (low), 30°C (high) |
| 2 | 0.3 mM (high), 15°C (low) | 0.3 mM (high), 30°C (high) |
| 3 | 0.03 mM (low), 30°C (high) | 0.3 mM (high), 15°C (low) |
| 4 | 0.03 mM (low), 15°C (low) | 0.3 mM (high), 30°C (high) |
| 5 | 0.03 mM (low), 15°C (low) | 0.3 mM (high), 15°C (low) |
| 6 | 0.03 mM (low), 30°C (high) | 0.3 mM (high), 30°C (high) |

The two fluids used in each Test represented different combinations of a low (0.03 mM) and high (0.3 mM) concentration of quinine, and lower (15°C) and higher (30°C) solution temperatures. In one Test, the low quinine concentration was proffered at a low (fluid 1) and high (fluid 2) temperature across brief-access trials (i.e., only temperature varied; **Table 1**, Test 1). Another Test repeated this procedure but using the high quinine concentration (Test 2). Two additional Tests proffered mice both quinine concentrations with temperature varying inversely with concentration (i.e., the low concentration was 30°C, high 15°C; Test 3) or in parallel (i.e., the low concentration was 15°C, high 30°C; Test 4). Finally, two Tests presented both the low and high concentration of quinine at a constant lower (Test 5) or higher (Test 6) fluid temperature (i.e., only concentration varied). These conditions were selected to investigate additive effects of concentration and temperature on quinine responses. During each Test, the two solutions were individually offered on 10-sec trials, with solution order randomized within each of 10 contiguous trial blocks (20 trials total; mice were allowed up to 100 sec of licking access to each solution per test day). Each quinine Test was presented once to each mouse, for a total of six brief-access sessions.

#### 2.6.2. Sucrose and glucose experiments

Two squads of mice were tested with sucrose and two additional squads with glucose; mice received either sucrose or glucose, but not both. Brief-access fluid exposure procedures with sugars were performed in two phases. Phase 1 was an orientation session to allow mice to learn that the sipper tubes offered sugar solutions, which are naturally appetitive to rodents. To do this, overnight water-restricted mice were proffered room temperature solutions of 300 mM sucrose or glucose in brief-access exposure sessions conducted over 2 consecutive days. During each session, mice were allowed to lick the sugar fluid over 20, 10-sec long trials.

Once phase 1 completed, mice were returned to their home cage and given *ad libitum* access to water. They then entered phase 2, which began the next day. Phase 2 monitored brief-access licking responses to temperature-controlled solutions of sucrose or glucose in water-replete (not thirst-motivated) mice with continuous access to water in their home cage. Mice also had free access to food in their home cage, so it was presumed that the preferred hedonic tone of the sugars, but not thirst or hunger, was the primary motivator of licking in phase 2.

Each Test offered mice two temperature-controlled solutions of one sugar. The solutions represented different combinations of low (100 mM) and high (500 mM) sugar concentrations and lower (15°C) and higher (30°C) solution temperatures (**Table 2**). These conditions were analogous to those used for quinine and included Tests where only temperature (Tests 1 and 2) or concentration (Tests 5 and 6) differed, and Tests where concentration and temperature varied inversely (Test 3) or in parallel (Test 4). Sugar solutions were singly presented in discrete 10-sec trials, with solution order randomized within 10 contiguous trial blocks (20 trials total; mice were allowed up to 100 sec of licking access to each solution per day). Each sugar Test condition was repeated over three consecutive days in a Test Block, for a total of 18 brief-access test sessions across the 6 Blocks. This aimed to allow sufficient data collection using non-thirst-motivated mice. Test condition order was randomized for each mouse using a Latin square design to control for potential longitudinal effects of exposure.

**Table 2.**
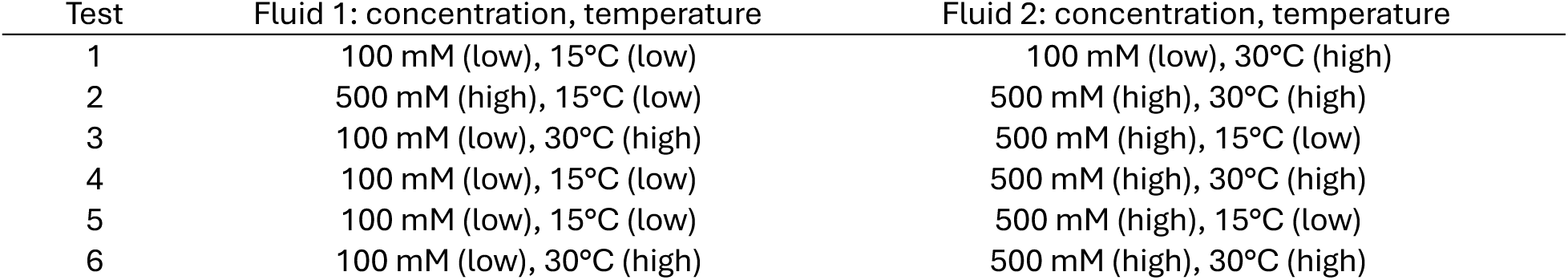
Stimulus concentrations and temperatures used in two-fluid brief-access thermolickometry tests with sugars (sucrose or glucose).

| Test | Fluid 1: concentration, temperature | Fluid 2: concentration, temperature |
|---|---|---|
| 1 | 100 mM (low), 15°C (low) | 100 mM (low), 30°C (high) |
| 2 | 500 mM (high), 15°C (low) | 500 mM (high), 30°C (high) |
| 3 | 100 mM (low), 30°C (high) | 500 mM (high), 15°C (low) |
| 4 | 100 mM (low), 15°C (low) | 500 mM (high), 30°C (high) |
| 5 | 100 mM (low), 15°C (low) | 500 mM (high), 15°C (low) |
| 6 | 100 mM (low), 30°C (high) | 500 mM (high), 30°C (high) |

The low and high concentrations of quinine and sugar we tested elicit differences in licking behavior in B6 mice (Ascencio Gutierrez et al., 2024; Boughter et al., 2005). The two fluid temperatures examined show dependence on TRPM8 for oral sensation. Thirty degrees Celsius is near the cooling activation threshold for TRPM8 while 15°C induces strong current flow through this ion channel (McKemy et al., 2002). Furthermore, these temperatures do not affect water licking in B6 mice but cause acceptance (15°C) and avoidance (30°C) of water in KO mice during brief-access tests conducted under water restriction, implying recognition of their orosensory valence relies on TRPM8 signaling (Li et al., 2024). Our thermolickometer was able to maintain these temperatures with similar precision across all trials analyzed (15°C set point: actual mean [± SD] fluid temperature = 15.1 ± 0.43°C; 30°C set point: actual mean fluid temperature = 30.0 ± 0.42°C).

### 2.7. Data analysis and statistics

The hypotheses and analytic plan underlying temperature, mouse line, and concentration effects were specified prior to data collection. It was anticipated that contextual and longitudinal effects would affect licking behavior, albeit not to the extent observed in this study. Additional post-hoc hypotheses regarding contextual and longitudinal effects on lick behavior were generated based on preliminary analysis of the results and subsequently analyzed and reported herein.

The number of licks mice emitted on each brief-access trial were calculated by adding 1 to the number of inter-lick intervals that were >50 msec. This criterion removed potential false licks attributable to electrical/RF noise. Trials with 1 or 2 licks were considered non-sampled and zeroed. The mean number of licks each mouse made to a stimulus per trial was then calculated. Mean licks per trial is referred to herein as “licks” for convenience.

Licks to quinine were standardized as water-adjusted licks (WAL) which were calculated by subtracting each mouse’s licks to room temperature water, calculated from the initial 5 trials of the two-day brief-access training period, from their licks to quinine. This expressed quinine responses for each mouse relative to an estimate of their water licking, with negative values reflecting the number of licks less than their water response (i.e., relative avoidance). Water was not included during quinine tests to force mice to direct their licks between two bitter solutions that differed in concentration and/or temperature in the context of solution palatability and their thirst. This strategy was adapted from single-bottle analyses of quinine licking in rodents (Hsiao & Fan, 1993; Spector & St John, 1998; Yamamoto et al., 1985).

#### 2.7.1. Model fitting

To analyze lick responses, two linear mixed models (LMMs) were fit, one to the quinine data and one to the sugar data. A random intercept at the mouse-level was included in all models. All subordinate interactions and main effects were included for each interaction. Model terms are referred to in italics below.

#### 2.7.2. Quinine model

Model fitting was conducted by evaluating all possible individual 2-, 3-, and 4-way interactions between temperature, concentration, mouse line, and sex; each of these variables’ 2-way interactions with exposure (defined below) were also evaluated, as well as all 2-way interactions between line, sex, and test. These candidate models were reviewed, prioritizing those with significant, biologically plausible interactions that improved model fit (as indicated by Akaike Information Criteria [AIC], log-likelihood, and residual error; **Table S1**). This produced the final model (**Table S2**; **Table S1**: M2o) in **Equation 1**.

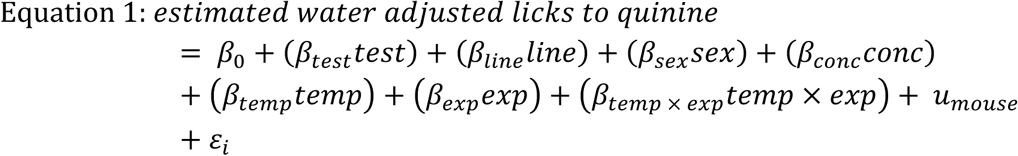

The variable *test* represents which of the 6 Tests an observation was obtained from (**Table 1**), *line* represents the observation’s mouse line (B6 or KO), *sex* represents the sex of the mouse (female or male), *conc* represents solution concentration [quinine] (0.03 mM or 0.3 mM), *temp* represents solution temperature (15°C or 30°C), and *exp* represents the number of previous exposures to the experimental procedure (i.e. days where the mouse was presented any thermal-taste solutions). To account for repeated measures and within-mouse variability, a random intercept for mouse (*u_mouse_*) was included. *ε_i_* represents the residual error. All variables were categorical except *exp* (exposure), which was continuous. Coefficient estimates are given in **Table S2**.

### 2.7.3. Sugar model

Model fitting was conducted by evaluating several structural and scientific LMMs derived from our experimental design and hypotheses (**Table S3**). Initially, we evaluated models using the raw lick means but found that their residuals possessed substantial heteroscedasticity due to a strongly right-skewed distribution of lick data. To approximate a normal distribution, we passed the sugar lick data through a log+1 transformation via MATLAB’s *log1p()* function and repeated our model evaluation. Unless otherwise noted, all model outcomes were back transformed into true lick units via exponentation (**Equation 2**) via MATLAB’s *expm1()* function prior to reporting and visualization herein.

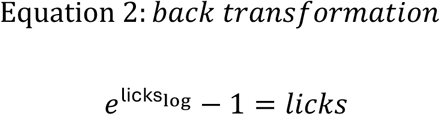

Model evaluation was conducted by assessing improvements in model fit parameters, conducting likelihood-ratio tests to its respective reference model(s), and general scientific plausibility/relevance. First, we evaluated a model that only included terms for structural, longitudinal, and between-mouse factors (M0) to validate the data structure (**Table S3**). Then we refit this model with the core scientific terms of *line*, *temp*, and *conc* and their 2- and 3-way interactions to serve as a general reference model (M1). Next, we evaluated a set of prespecified candidate models that explored various interactions and effects related to our hypotheses: longitudinal effects (M2), within-mouse interactions between *line* and *sex* (M3), and a structural evaluation of a potential *blockDay-test* interaction (M4; see below). After observing significant and near-significant interactions between *exposure* and *line* (M2a) and *conc* (M2c), we evaluated two additional models to determine if these effects interacted (M2f-g). Based on model fit criteria, we evaluated M2g’s adequacy as a potential final analytical model. We observed a modestly left-skewed residual distribution, though this was greatly reduced from the original untransformed lick model. No convergence issues or boundary estimates were encountered, and we selected M2g as the final analytical sugar model as in **Equation 3**.

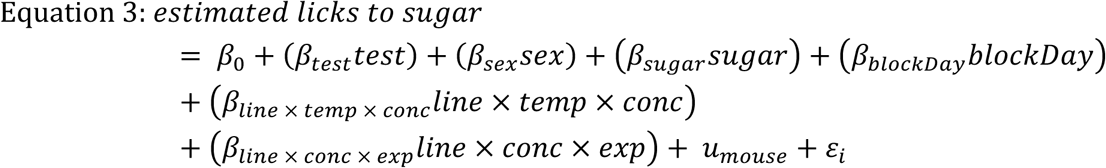

The variable *test* represents which of the 6 Tests an observation was obtained from (**Table 2**), *line* represents the observation’s mouse line (B6 or KO), *sex* represents the sex of the mouse (female or male), *conc* represents concentration [sugar] (100 mM or 500 mM), *temp* represents sugar fluid temperature (15°C or 30°C), *sugar* represents which sugar solution was used (glucose or sucrose), and *exp* represents the number of previous exposures to the experimental procedure. Unlike quinine, where each mouse was exposed to each test only once, the sugar protocol presented each test three times in a continuous block. To account for this in the model, we included *blockDay*, which represents the number of previous exposures to a given test (0, 1, or 2). A random intercept for mouse (*u_mouse_*) was included to account for repeated measures and within-mouse variability. *ε_i_* represents the residual error. A significant 3-way interaction between *line*, *conc*, and *exp* was included (**Table S3**: M2g).

Mice were run in four squads of 12, and each squad received only one sugar (2 squads per *sugar*). *Squad* was evaluated as a potential fixed or random effect, but was ultimately omitted due to, respectively, a lack of improvement in likelihood and near-zero estimated variance component (i.e., a boundary estimate). Since *squad* was perfectly collinear with *sugar*, and since *sugar* was more relevant to our experimental design, we opted to include *sugar* as the structural term instead. All variables were categorical except *exp*, which was continuous. Coefficient estimates are given in **Table S4**.

#### 2.7.4. Estimated marginal means, contrasts, and error terms

For both sugar and quinine data, the primary method of analysis was conducting contrasts between non-conditional (fixed effects) estimated marginal means (EMMs) generated by the final analytical models. First, a granular prediction grid design matrix was constructed, consisting of row vectors defining every valid condition an observation was collected from (quinine: 288, sugar: 1728). This matrix was then collapsed (averaged) into EMM coefficient matrices as required to conduct individual analyses. EMMs were generated by multiplying the EMM coefficient matrix by its respective model’s coefficient vector.

For quinine, EMM standard errors and 95% confidence intervals (CIs) were derived from the model’s variance-covariance matrix.

For log-transformed sugar data, point estimates (such as EMMs and their contrasts) were generated in the model’s native log-scale, then back-transformed into true lick units via exponentiation (**Equation 2**). To adjust for log-scale’s non-linearity, the average of the model’s error term and random-effect term were added to the log-scale point estimates immediately prior to back-transformation (Ramirez-Aldana & Naranjo, 2021, equation 5). For calculating CIs, we adapted the Krinsky and Robb method (Haab & McConnell, 2002, pg 112). Briefly, we used MATLAB’s *mvnrnd()* function to generate 1000 simulated coefficient vectors from the model’s coefficient variance-covariance matrix, which were then multiplied against the prediction grid design matrix to produce 1000 simulated estimates for each of the 1728 conditions. These log-scale estimates were then back-transformed into true lick units and sorted numerically. For a given condition, the lower and upper 95% CI bounds were defined as the 2.5^th^ and 97.5^th^ percentiles, respectively, of its simulated estimates. Standard errors for individual estimates were also derived from these sets of simulated estimates, by taking the standard deviation of their sampling distribution after back-transformation.

In the longitudinal analyses, slopes were defined as the EMM of *temp* (for quinine) or *line:conc* (for sugar) at *exp n+1* minus the EMM at *exp n* (i.e. the change associated with a 1-unit increase in *exp*). For sugar, due to the non-linear back-transformation of the model’s log-scale estimates to true licks units, the slope values varied depending on which specific *exps* were contrasted. To account for this, we took the difference between the *line:conc* EMMs at *exp 0* and *exp 15* (i.e. total change across the study period), then divided this value by 15 to produce an average slope in units ‘change in licks per 1-unit increase in *exp*’.

All analyses, estimates, and contrasts were conducted in MATLAB R2025a (The MathWorks, Natick, MA). Models were fit using the maximum likelihood method via *fitlme()*. Contrasts were conducted using *coefTest()* with residual degrees of freedom (DF_2_ = number of observations - number of fixed effects). The α was set to 0.05. A false-discovery rate (FDR) correction was applied to the *p*-values to adjust for multiple comparisons (Benjamini & Hochberg, 1995).

## 3. Results

### 3.1. Quinine

Using brief-access thermolickometry, we measured orosensory responses to pairs of temperature- and concentration-controlled quinine solutions from 12 B6 (6 female) and 10 KO (4 female) mice. Mice were water-restricted for all quinine experiments.

#### 3.1.1. Concentration and temperature independently and summatively modulate licking behavior to quinine

We first investigated the effect of acceptance to quinine solutions when their temperature (either 15°C or 30°C) or concentration (either 0.03 mM or 0.3 mM) was varied within the same Test (**Table 1**). Given that we observed no significant interaction between concentration and temperature in the quinine model (**Table S1**: M2f), we collapsed the data across concentration for Tests 1-2 (**Figure 1A**, **Table 3 A-B**) and across temperature for Tests 5-6 (**Figure 1B**, **Table 3 C-D**). Tests 3 and 4 evaluated inverted (**Figure 2A**, **Table 3 E**) and parallel (**Figure 2B**, **Table 3 F**) changes, respectively, in concentration and temperature to evaluate the relative magnitude and potential summation of their effects (**Table 3G**). In addition, we collapsed across *line* and *sex* for the above-described comparisons due to a lack of significant interactions with *conc* and *temp*.

**Figure 1.**
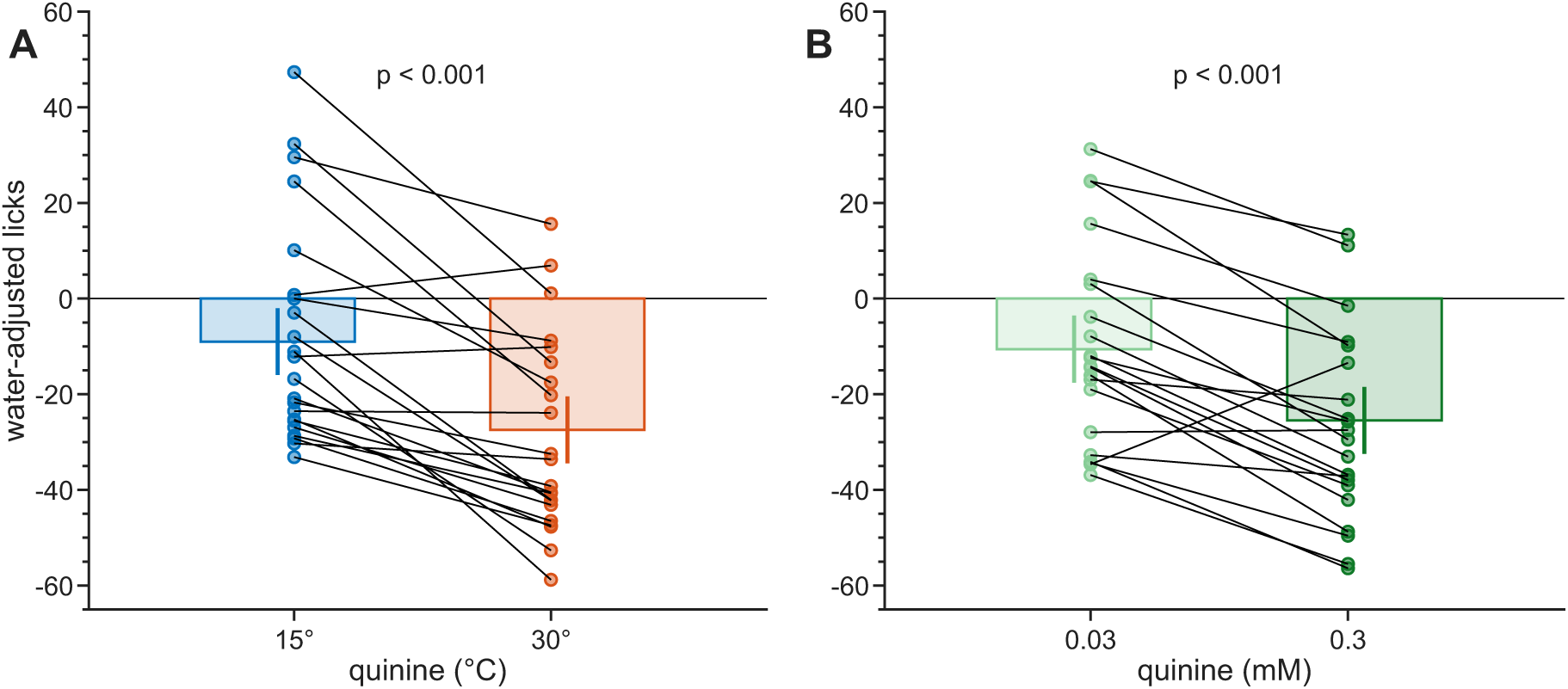
Lick responses to quinine solutions presented at different temperatures and constant concentrations (A) and different concentrations at constant temperatures (B). Due to a lack of significant interactions, data were collapsed across line and concentration in (A), and across line and temperature in (B). The observed mean water-adjusted licks per solution are plotted as circles, one for each mouse; solid lines connect observations from the same mouse between the two solutions. The estimated marginal means derived from these data are overlaid as columns. Error bars indicate 95% confidence intervals. (A) Results from Tests 1 and 2 (Table 1), showing responses to two different temperatures (15°C and 30°C) when presented together at the same concentration. (B) Results from Tests 5 and 6 (Table 1), showing responses to two different concentrations (0.03 mM or 0.3 mM) when presented together at the same temperature. Significant effects of concentration and temperature were observed (p < 0.001).

##### 3.1.1.1. Only temperature varies

EMMs and contrasts revealed a significant effect of temperature (**Figure 1A**; **Table 1**, Tests 1-2), whereby mice emitted 18.44 [14.55, 22.34] fewer estimated water-adjusted licks (WAL) to 30°C quinine than 15°C quinine (**Table 3 A-B**; *F*_(1,250)_ = 86.87, *p* = 1.1 × 10^-17^). Within these data, responses did not significantly vary by quinine concentration: 0.3 mM received 5.8 [0.3, 11.3] fewer WAL than 0.03 mM (*F_(_*_1,250)_ = 4.3, p = 0.0539).

##### 3.1.1.2. Only concentration varies

Similarly, we observed a significant effect of concentration (**Figure 1B**; **Table 1**, Tests 5-6), whereby 0.3 mM quinine evoked 14.92 [11.00, 18.84] fewer estimated WAL than 0.03 mM quinine (**Table 3 C-D**; *F*_(1,250)_ = 56.21, *p* = 1.3 × 10^-12^). Within these data, responses did not significantly vary by temperature: 30°C received 4.9 [-0.7, 10.5] fewer WAL than 15°C (*F*_(1,250)_ = 3.0, *p* = 0.0855).

##### 3.1.1.3. Both concentration and temperature vary

When solution concentration and temperature were inversely varied (**Table 1**, Test 3) – i.e., low-concentration at high-temperature (0.03 mM @ 30°C) alongside high-concentration at lower-temperature (0.3 mM @ 15°C) – mice emitted similar WAL to each quinine solution (**Figure 2A**; **Table 3E**). Specifically, mice made 3.53 [-2.00, 9.05] fewer WAL to the ‘low concentration (conc) – high temperature (temp)’ 0.03 mM @ 30°C solution than the ‘high conc – low temp’ 0.3 mM @ 15°C solution; this contrast was not significant (*F*_(1,250)_ = 1.58, *p* = 0.2101). However, when low-concentration low-temperature (0.03 mM @ 15°C) and high-concentration high-temperature (0.3 mM @ 30°C) quinine solutions were jointly presented (**Table 1**, Test 4), we observed that mice emitted 33.36 [27.83, 38.89] fewer WAL to the ‘high conc - high temp’ 0.3 mM @ 30°C than to the ‘low conc - low temp’ 0.03 mM @ 15°C solution (**Figure 2B**; **Table 3 F**; *F*_(1,250)_ = 141.35, *p* = 2.7 × 10^-25^). A second-order contrast between these ‘inverse’ and ‘parallel’ contrasts (**Table 3 G**) found that parallel summation of concentration and temperature resulted in a significantly larger difference than their inverse (*F*_(1,250)_ = 86.87, *p* = 1.1 × 10^-17^).

**Table 3.** Estimated marginal means for each quinine Test (Table 1), averaged across all other factors, and their associated contrasts and test statistics. Contrasts were conducted between the four quinine solutions presented in Tests 1 + 2 (collapsed across concentration), the four quinine solutions presented in Tests 5 + 6 (collapsed across temperature), the two quinine solutions presented in Test 3, and the two quinine solutions presented in Test 4. An additional second-order contrast evaluated the between-contrast difference for E and F (Tests 3 and 4). CI = 95% Confidence Interval, conc = concentration, temp = temperature. * = p < 0.05. Note, Contrast Estimate = Solution 1 Estimate -Solution 2 Estimate.

| ID | Test | Contrast Description | Solution 1 | Solution 1 Estimate [CI] | Solution 2 | Solution 2 Estimate [CI] | Contrast Estimate [CI] | F<sub>(1,250)</sub> | p-value |
|---|---|---|---|---|---|---|---|---|---|
| A | 1 | Between *temp* | 0.03 mM @ 30°C | -24.54<br>[-32.04, -17.04] | 0.03mM @ 15°C | -6.10<br>[-13.60, 1.41] | -18.44<br>[-22.34, -14.55] | 86.87 | 1.1x10<sup>-17</sup> * |
| B | 2 | Between *temp* | 0.3 mM @ 30°C | -30.36<br>[-37.86, -22.86] | 0.3 mM @ 15°C | -11.92<br>[-19.42, -4.41] |  |  |  |
| C | 5 | Between *conc* | 0.3 mM @ 15°C | -23.04<br>[-30.61, -15.47] | 0.03 mM @ 15°C | -8.13<br>[-15.70, -0.56] | -14.92<br>[-18.84, -11.00] | 56.21 | 1.3x10<sup>-12</sup> * |
| D | 6 | Between *conc* | 0.3 mM @ 30°C | -27.95<br>[-35.45, -20.44] | 0.03 mM @ 30°C | -13.03<br>[-20.54, -5.52] |  |  |  |
| E | 3 | Between *conc:temp* | 0.3 mM @ 15°C | -17.22<br>[-24.98, -9.47] | 0.03 mM @ 30°C | -20.75<br>[-28.51,-13.00] | 3.53<br>[-2.00, 9.05] | 1.58 | 0.2101 |
| F | 4 | Between *conc:temp* | 0.3 mM @ 30°C | -35.95<br>[-43.71, -28.20] | 0.03 mM @ 15°C | -2.59<br>[-10.35, 5.16] | -33.36<br>[-38.89, -27.83] | 141.35 | 2.7x10<sup>-25</sup> * |
| G | 3 v 4 | Between Contrast | Test 4 Contrast | -33.36<br>[-38.89, -27.83] | Test 3 Contrast | 3.53<br>[-2.00, 9.05] | -36.89<br>[-44.68, -29.09] | 86.87 | 1.1x10<sup>-17</sup> * |

**Figure 2.**
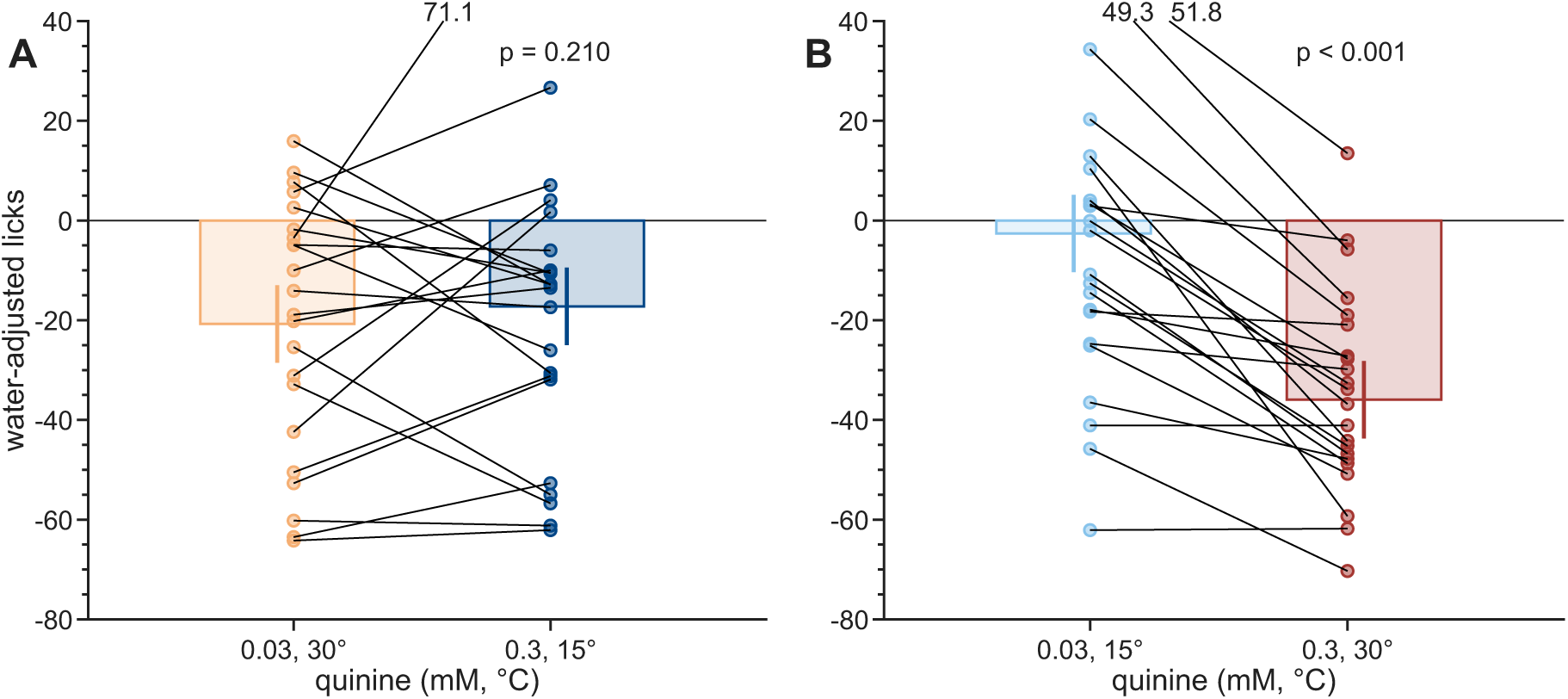
Lick responses to quinine solutions of different temperatures and concentrations. Due to the lack of a significant interaction, data were collapsed across mouse line. The observed mean water-adjusted licks per solution are plotted as circles, one for each mouse; lines connect observations from the same mouse between the two solutions. Real numbers at the top of the plots denote actual values for points above the Y-axis scale. The estimated marginal means derived from these data are overlaid as columns. Error bars indicate 95% confidence intervals. (A) Results from Test 3 (Table 1), showing responses to solutions with inversely varied concentrations and temperatures. (B) Results from Test 4 (Table 1), showing responses to solutions where concentration and temperature varied in parallel.

Altogether, these data suggest that lower temperature and lower concentration both independently increase acceptance of quinine solutions (**Figure 1**; **Table 3**). The effects of concentration and temperature are proportional in size and summative, as evinced by their very low contrast when inversely varied (**Figure 2A**) and very high contrast when varied in parallel (**Figure 2B**).

#### 3.1.2. Acceptance of quinine solutions is context-dependent

Next, we asked if mice would emit the same number of licks to the same quinine solution presented in different experimental contexts (i.e., Tests). Specifically, when presented alongside quinine solutions of different concentrations and/or temperatures. Our experimental design consisted of four solutions with unique concentration-temperature properties: 0.03 mM @ 15°C, 0.03 mM @ 30°C, 0.3 mM @ 15°C, and 0.3 mM @ 30°C. Every solution was presented in three Tests, each time paired with one of the other three solutions (**Table 1**). This allowed us to generate EMM WAL to a given ‘reference solution’ in each of its three experimental contexts (**Table S5**), which we then contrasted to determine if responses to the reference solution significantly varied based on the context solution it was provided alongside. The specific contrasts referenced below for quinine are detailed in **Table 4** and **Figure 3**.

Mice emitted -6.13 [-17.78, 5.52] WAL to **0.03 mM @ 15°C** when it was presented alongside the higher-temperature 0.03 mM @ 30°C, -8.16 [-19.85, 3.53] WAL when tested with the higher-concentration 0.3 mM @ 15°C, and -2.63 [-14.44, 9.19] WAL when presented with the higher-concentration higher-temperature 0.3 mM @ 30°C. No significant differences were observed between these estimates (*F*_(1,250)_ ≤ 3.39, *p* ≥ 0.1126; Contrasts A-C; **Figure 3A**).

Mice emitted -24.56 [-36.21, -12.91] WAL to **0.03 mM @ 30°C** when it was presented alongside the lower-temperature 0.03 mM @ 15°C context solution, -20.77 [-32.60, -8.96] WAL when presented alongside the higher-concentration lower-temperature 0.3 mM @ 15°C, and -13.05 [-24.71, -1.40] WAL when presented alongside the higher-concentration 0.3 mM @ 30°C. Mice emitted significantly fewer WAL to **0.03 mM @ 30°C** when it was presented alongside 0.03 mM @ 15°C (*F*_(1,250)_ = 13.51, *p* = 0.0032; Contrast D; **Figure 3B**) or 0.3 mM @ 15°C (*F*_(1,250)_ = 6.77, *p* = 0.0295; Contrast F; **Figure 3B**) than with 0.3 mM @ 30°C, suggesting the availability of lower temperature quinine fluids lessen its appeal. No significant difference was observed between the two 15°C contexts (*F*_(1,250)_ = 1.63, *p* = 0.2710; Contrast E; **Figure 3B**).

**Table 4.**
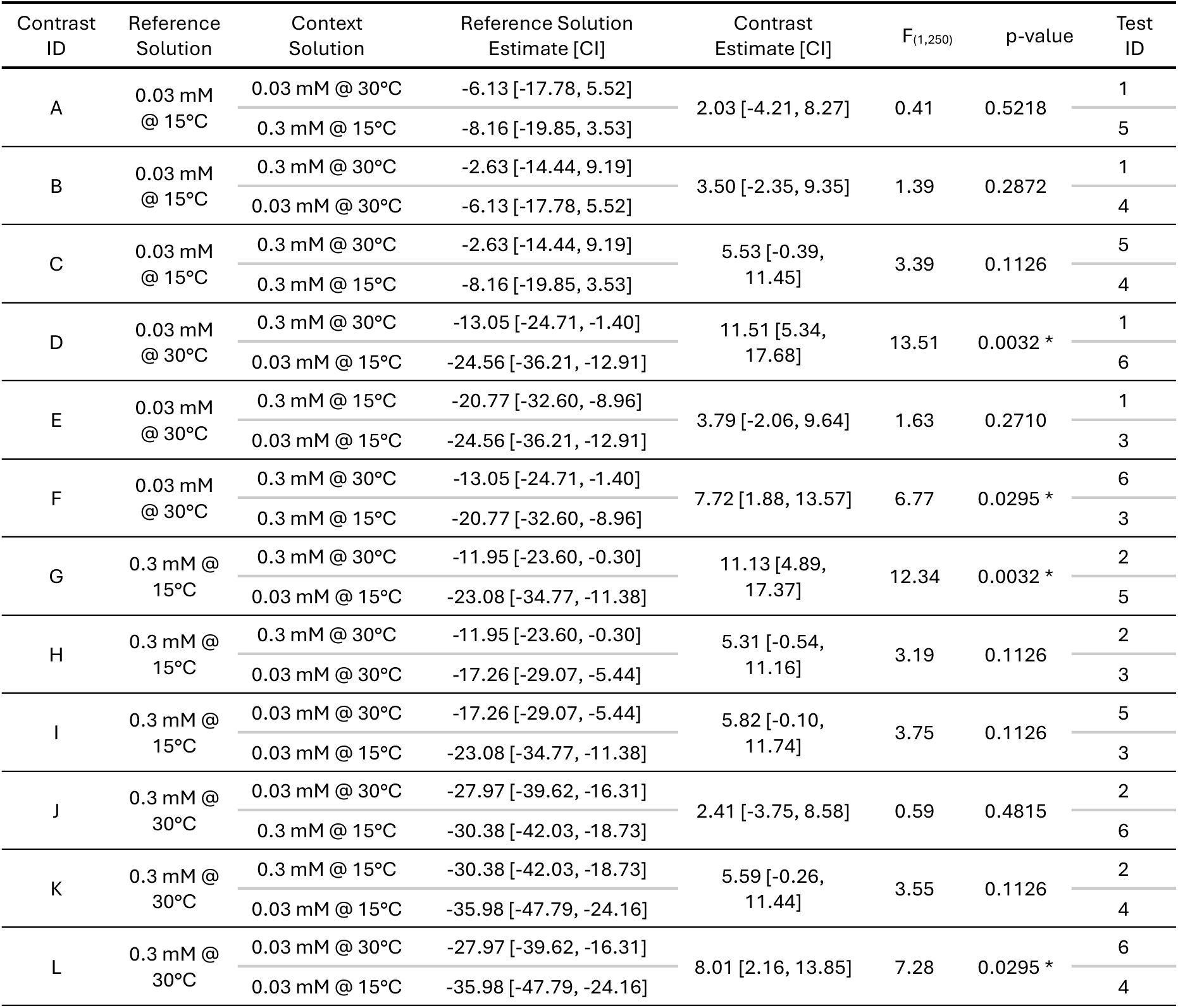
Outcomes of contrasts between the estimated marginal mean water-adjusted licks for a given reference solution when paired alongside each of the other three context solutions as indicated by its Test ID (see Table 1). Each row reports the estimated marginal mean [95% confidence interval] water-adjusted licks to a given reference solution when it was presented alongside one of its three possible context solutions. Pairwise contrasts were conducted between each of the possible pairings of a given reference solution’s three context solutions . CI = 95% Confidence Interval, * = p < 0.05. Note, Contrast Estimate = First Reference Solution Estimate - Second Reference Solution Estimate. A higher Contrast Estimate value indicates greater aversion to the Second Reference Solution.

| Contrast ID | Reference Solution | Context Solution | Reference Solution Estimate [CI] | Contrast Estimate [CI] | F<sub>(1,250)</sub> | p-value | Test ID |
|---|---|---|---|---|---|---|---|
| A | 0.03 mM @ 15°C | 0.03 mM @ 30°C | -6.13 [-17.78, 5.52] | 2.03 [-4.21, 8.27] | 0.41 | 0.5218 | 1 |
|  |  | 0.3 mM @ 15°C | -8.16 [-19.85, 3.53] |  |  |  | 5 |
| B | 0.03 mM @ 15°C | 0.3 mM @ 30°C | -2.63 [-14.44, 9.19] | 3.50 [-2.35, 9.35] | 1.39 | 0.2872 | 1 |
|  |  | 0.03 mM @ 30°C | -6.13 [-17.78, 5.52] |  |  |  | 4 |
| C | 0.03 mM @ 15°C | 0.3 mM @ 30°C | -2.63 [-14.44, 9.19] | 5.53 [-0.39, 11.45] | 3.39 | 0.1126 | 5 |
|  |  | 0.3 mM @ 15°C | -8.16 [-19.85, 3.53] |  |  |  | 4 |
| D | 0.03 mM @ 30°C | 0.3 mM @ 30°C | -13.05 [-24.71, -1.40] | 11.51 [5.34, 17.68] | 13.51 | 0.0032 * | 1 |
|  |  | 0.03 mM @ 15°C | -24.56 [-36.21, -12.91] |  |  |  | 6 |
| E | 0.03 mM @ 30°C | 0.3 mM @ 15°C | -20.77 [-32.60, -8.96] | 3.79 [-2.06, 9.64] | 1.63 | 0.2710 | 1 |
|  |  | 0.03 mM @ 15°C | -24.56 [-36.21, -12.91] |  |  |  | 3 |
| F | 0.03 mM @ 30°C | 0.3 mM @ 30°C | -13.05 [-24.71, -1.40] | 7.72 [1.88, 13.57] | 6.77 | 0.0295 * | 6 |
|  |  | 0.3 mM @ 15°C | -20.77 [-32.60, -8.96] |  |  |  | 3 |
| G | 0.3 mM @ 15°C | 0.3 mM @ 30°C | -11.95 [-23.60, -0.30] | 11.13 [4.89, 17.37] | 12.34 | 0.0032 * | 2 |
|  |  | 0.03 mM @ 15°C | -23.08 [-34.77, -11.38] |  |  |  | 5 |
| H | 0.3 mM @ 15°C | 0.3 mM @ 30°C | -11.95 [-23.60, -0.30] | 5.31 [-0.54, 11.16] | 3.19 | 0.1126 | 2 |
|  |  | 0.03 mM @ 30°C | -17.26 [-29.07, -5.44] |  |  |  | 3 |
| I | 0.3 mM @ 15°C | 0.03 mM @ 30°C | -17.26 [-29.07, -5.44] | 5.82 [-0.10, 11.74] | 3.75 | 0.1126 | 5 |
|  |  | 0.03 mM @ 15°C | -23.08 [-34.77, -11.38] |  |  |  | 3 |
| J | 0.3 mM @ 30°C | 0.03 mM @ 30°C | -27.97 [-39.62, -16.31] | 2.41 [-3.75, 8.58] | 0.59 | 0.4815 | 2 |
|  |  | 0.3 mM @ 15°C | -30.38 [-42.03, -18.73] |  |  |  | 6 |
| K | 0.3 mM @ 30°C | 0.3 mM @ 15°C | -30.38 [-42.03, -18.73] | 5.59 [-0.26, 11.44] | 3.55 | 0.1126 | 2 |
|  |  | 0.03 mM @ 15°C | -35.98 [-47.79, -24.16] |  |  |  | 4 |
| L | 0.3 mM @ 30°C | 0.03 mM @ 30°C | -27.97 [-39.62, -16.31] | 8.01 [2.16, 13.85] | 7.28 | 0.0295 * | 6 |
|  |  | 0.03 mM @ 15°C | -35.98 [-47.79, -24.16] |  |  |  | 4 |

**Figure 3.**
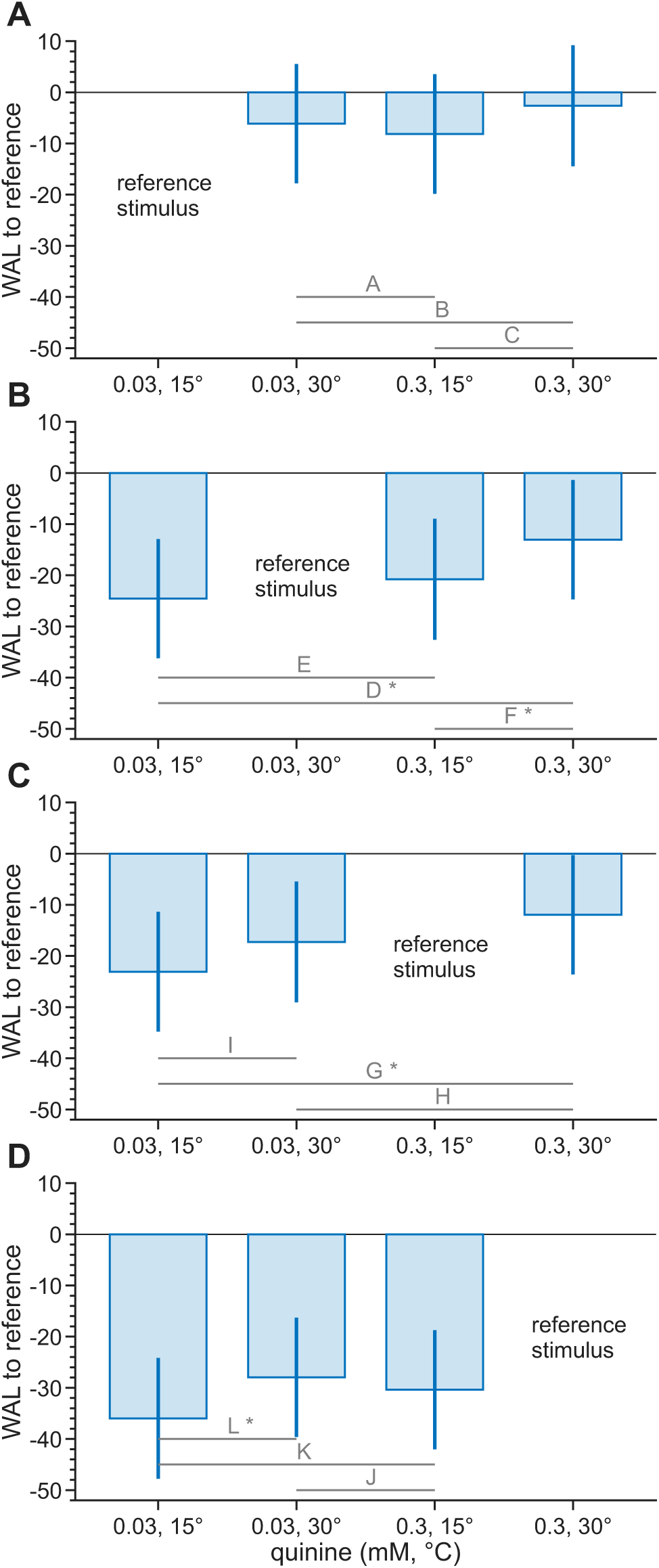
Estimated marginal means of licks to each reference quinine solution in its three contexts. Each subplot (A-D) corresponds to one of the four reference conditions. Bars show water-adjusted licks (WAL) to the reference solution (y-axes) when presented alongside each of the three context solutions (x-axes). Horizontal lines and letters denote specific contrast tests performed (Table 4). Error bars represent the 95% confidence interval. * = p < 0.05

Mice emitted -23.08 [-34.77, - 11.38] WAL to **0.3 mM @ 15°C** when it was presented alongside the lower-concentration 0.03 mM @ 15°C, -17.26 [-29.07, -5.44] WAL when presented alongside the lower-concentration higher-temperature 0.03 mM @ 30°C, and -11.95 [-23.60, -0.30] WAL when presented alongside the higher-temperature 0.3 mM @ 30°C. Mice emitted significantly fewer WAL to **0.3 mM @ 15°C** when it was presented alongside 0.03 mM @ 15°C than with 0.3 mM @ 30°C (*F*_(1,250)_ = 12.34, *p* = 0.0032; Contrast G; **Figure 3C**). No significant difference was observed between the 0.03 mM @ 30°C and 0.3 mM @ 30°C contexts (*F*_(1,250)_ = 3.19, *p* = 0.1126; Contrast H; **Figure 3C**) or between the 0.03 mM @ 30°C and 0.03 mM @ 15°C contexts (*F*_(1,250)_ = 3.75, *p* = 0.1126; Contrast I; **Figure 3C**).

Finally, mice emitted -35.98 [-47.79, -24.16] WAL to **0.3 mM @ 30°C** when it was presented alongside the lower-concentration lower-temperature 0.03 mM @ 15°C, -27.97 [-39.62, -16.31] WAL when presented alongside the lower-concentration 0.03 mM @ 30°C, and -30.38 [-42.03, -18.73] WAL when tested with the lower-temperature 0.3 mM @ 15°C. Mice emitted significantly fewer WAL to **0.3 mM @ 30°C** when it was presented alongside 0.03 mM @ 15°C than alongside 0.03 mM @ 30°C (*F*_(1,250)_ = 7.28, *p* = 0.0295; Contrast L; **Figure 3D**), suggesting that the availability a cooler, lower concentration quinine solution reduces its acceptance. No significant difference was observed between the 0.03 mM @ 30°C and 0.3 mM @ 15°C contexts (*F*_(1,250)_ = 0.59, *p* = 0.4815; Contrast J; **Figure 3D**) or between the 0.3 mM @ 15°C and 0.03 mM @ 15°C contexts (*F*_(1,250)_ = 3.55, *p* = 0.1126; Contrast K; **Figure 3D**).

Altogether, our observations indicate that licking activity to quinine is context dependent. Analysis suggests that a difference in temperature between the context solutions is the main factor driving changes in licking to the reference solution; a similar but non-significant trend was also observed in concentration. The largest effect sizes (i.e. EMM contrasts) were observed when both temperature and concentration varied in parallel (i.e. both increased or both decreased) (≥ 11.13 WAL; Contrasts D and G), then when only temperature varied (5.53 - 8.01 WAL; Contrasts C, F, I, and L), then when only concentration varied (3.50 - 5.59 WAL; Contrasts B, E, H, and K), and the smallest effect occurred when both concentration and temperature varied inversely (i.e. one increased while the other decreased) (≤ 2.41 WAL; Contrasts A and J). These observations mirror those presented in **Figures 1-2**, which indicate that both concentration and temperature can influence licks to quinine solutions.

#### 3.1.3. Experimental exposures are associated with increased licks to quinine in a temperature-dependent manner

During model fitting, we observed a significant longitudinal effect on licking behavior across the study period, whereby estimated WAL increased with every additional day of testing (‘exposure’) after accounting for other factors. In addition, our quinine LMM revealed a significant interaction between temperature and exposure (**Table S2**). Here, we explored this longitudinal effect and its potential temperature-dependency. Contrasts below are found in **Table 5** and depicted in **Figure 4**.

**Table 5.** Nine pair-wise contrasts were conducted between the estimated marginal means of water-adjusted licks in various temperature-exposure (*temp-exp*) conditions. Contrasts A-F compare 15°C and 30°C on a given *exp*. Contrasts G and H compare the first and second *exps* at each *temp* to estimate a slope. Contrast I compares Contrasts G and H to evaluate the interaction effect: i.e. is the effect of *exp* the same at 15°C and at 30°C? For each Contrast, the estimated marginal means, their estimated contrasts, and the F-statistic and FDR-adjusted p-value are reported. CI = 95% Confidence Interval, *exp* = exposure, *temp* = temperature, * = p < 0.05

| Contrast ID | Contrast Description | Condition | Estimated Mean [CI] | Estimated Contrast [CI] | F<sub>(1,250)</sub> | p-value |
|---|---|---|---|---|---|---|
| A | Between *temp*, *exp* 1 | 15°C, *exp* 1 | -33.83 [-41.16, -26.49] | 7.36 [1.55, 13.17] | 6.22 | 0.0133 * |
|  |  | 30°C, *exp* 1 | -41.18 [-48.51, -33.86] |  |  |  |
| B | Between *temp*, *exp* 2 | 15°C, *exp* 2 | -24.90 [-31.72, -18.08] | 9.99 [5.67, 14.31] | 20.72 | 1.07x10<sup>-5</sup> * |
|  |  | 30°C, *exp* 2 | -34.88 [-41.70, -28.06] |  |  |  |
| C | Between *temp*, *exp* 3 | 15°C, *exp* 3 | -15.97 [-22.52, -9.41] | 12.62 [9.28, 15.93] | 55.39 | 2.9x10<sup>-12</sup> * |
|  |  | 30°C, *exp* 3 | -28.58 [-35.13, -22.03] |  |  |  |
| D | Between *temp*, *exp* 4 | 15°C, *exp* 4 | -7.03 [-13.59, -0.48] | 15.24 [11.90, 18.59] | 80.65 | 2.0x10<sup>-16</sup> * |
|  |  | 30°C, *exp* 4 | -22.28 [-28.82, -15.74] |  |  |  |
| E | Between *temp*, *exp* 5 | 15°C, *exp* 5 | 1.90 [-4.94, 8.73] | 17.87 [13.54, 22.20] | 66.04 | 4.6x10<sup>-14</sup> * |
|  |  | 30°C, *exp* 5 | -15.98 [-22.77, -9.17] |  |  |  |
| F | Between *temp*, *exp* 6 | 15°C, *exp* 6 | 10.83 [3.47, 18.19] | 20.50 [14.68, 26.32] | 48.09 | 5.2x10<sup>-11</sup> * |
|  |  | 30°C, *exp* 6 | -9.68 [-16.99, -2.36] |  |  |  |
| G | *exp* Slope, 15°C | 15°C, *exp* 2 | -24.90 [-31.72, -18.08] | 8.93 [7.58, 10.29] | 168.53 | 7.6x10<sup>-29</sup> * |
|  |  | 15°C, *exp* 1 | -33.83 [-41.16, -26.49] |  |  |  |
| H | *exp* Slope, 30°C | 30°C, *exp* 2 | -34.88 [-41.70, -28.06] | 6.30 [4.96, 7.64] | 85.90 | 4.1x10<sup>-17</sup> * |
|  |  | 30°C, *exp* 1 | -41.18 [-48.51, -33.86] |  |  |  |
| I | Interaction, G v H | 15°C, *exp* 1 - 15°C, *exp* 2 | 8.93 [7.58, 10.29] | 2.63 [0.69, 4.57] | 7.10 | 0.0092 * |
|  |  | 30°C, *exp* 1 - 30°C, *exp* 2 | 6.30 [4.96, 7.64] |  |  |  |

**Figure 4.**
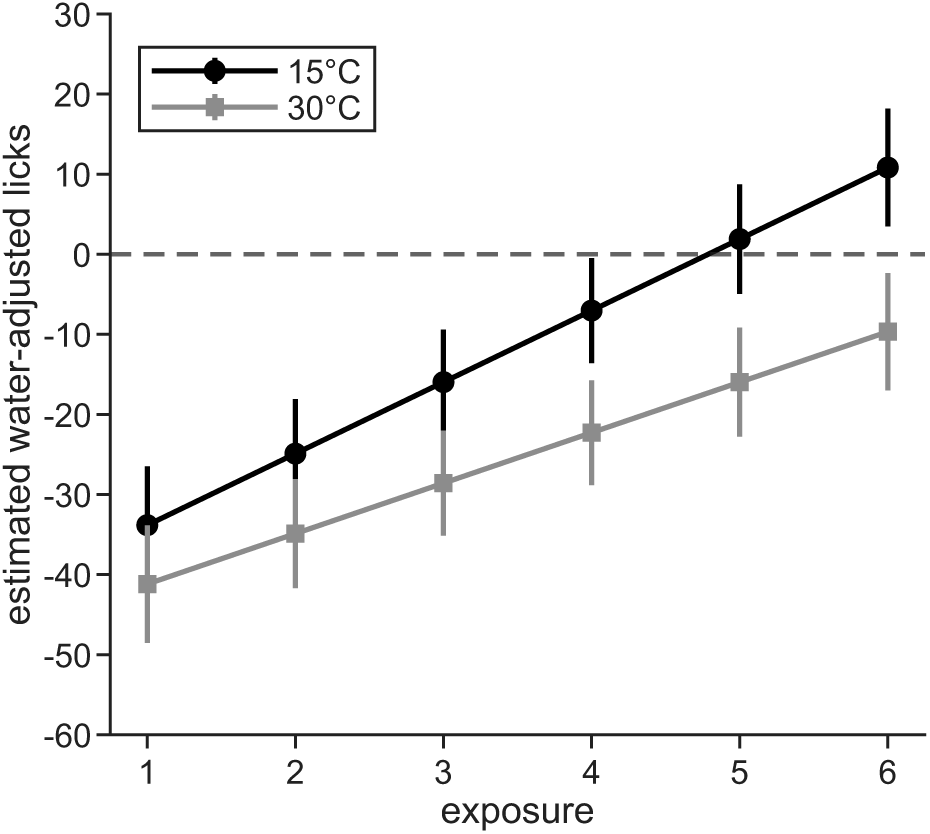
Estimated marginal means of water-adjusted licks (WAL) for each quinine exposure, plotted by-temperature. A lower estimated value indicates greater aversion to quinine. For every exposure, 15°C had a significantly higher estimated WAL than 30°C (p ≤ 0.0133). Moreover, the slope of 15°C is significantly higher than the slope of 30°C (p = 0.0092). Error bars represent the 95% confidence intervals. The dotted line at 0 represents an equal number of estimated licks to water and quinine.

We quantified the longitudinal effect of exposure (*exp*) at 15°C and 30°C using EMMs. At 15°C, we found that WAL to quinine increased at an estimated rate of 8.93 [7.58, 10.29] per exposure (*F*_(1,250)_ = 168.53, *p* = 7.6 × 10^-29^; Contrast G). At 30°C, a lower but still positive rate of 6.30 [4.96, 7.64] WAL per exposure was observed (*F*_(1,250)_ = 85.90, *p* = 4.1 × 10^-17^; Contrast H), with a significant difference of 2.63 [0.69, 4.57] between the two temperatures (*F*_(1,250)_ = 7.10, *p* = 0.0092; Contrast I). The between-temperature contrast increased from 7.36 [1.55, 13.17] WAL on the first exposure to 20.50 [14.68, 26.32] on the sixth and final exposure (Contrast A-F; *F*_(1,250)_ ≥ 6.22, *p* ≤ 0.0133). These outcomes suggest that a longitudinal effect of higher lick rate exists in our dataset, and that it increases significantly faster at 15°C than 30°C.

### 3.2. Sugar

Using brief-access thermolickometry, we measured orosensory responses to temperature- and concentration-controlled sucrose and glucose solutions from 23 B6 (11 female) and 24 KO (13 female) mice. Mice were allowed *ad libitum* access to water throughout testing to isolate the effect of thirst pressure from their acceptance of sugar solutions. To eliminate carryover effects, a given mouse was tested with only sucrose or glucose and was not included in the brief-access tests with quinine. Since we observed no significant difference in responses between sucrose and glucose (*F*_(1,1671)_ = 0.06, *p* = 0.8009), data were collapsed across both sugars for all analyses.

#### 3.2.1. Temperature and concentration affect licking to sugar solutions in a line-dependent manner

We first investigated the effect of acceptance to sugar solutions (glucose, sucrose) when their concentration (100 mM and 500 mM) and/or their temperature (15°C and 30°C) were varied. Line differences were evaluated because of a significant effect of *line* in the model.

##### 3.2.1.1. Only temperature varies

Contrasts between EMMs revealed a significant effect of temperature in KO, but not B6, mice (**Table 6 A-D**). KO mice emitted, on average, 1.10 [0.43, 1.90] more licks to 15°C than 30°C at 100 mM (**Table 2**, Test 1), and 1.45 [0.62, 2.45] more licks at 500 mM (**Table 2**, Test 2; *F*_(1,1671)_ ≥ 11.04, *p* ≤ 0.0009; **Figure 5A**). These effects were significant in the face of wide variability in individual responses, as most KO mice that avidly licked 15°C sugars decreased their licks to 30°C (**Figure 5A**). On the other hand, B6 mice emitted 0.15 [-0.58, 0.90] and 0.75 [-1.04, 2.70] more licks to 15°C than 30°C at 100 mM and 500 mM, respectively, and neither difference was significant (*F*_(1,1671)_ ≤ 0.67, *p* ≥ 0.4401; **Figure 5B**).

**Table 6.** Estimated marginal mean licks for each sugar Test condition, averaged across all other factors, and their associated contrasts and test statistics. Contrasts A-L were conducted between the two sugar solutions presented in Tests 1-6 (Table 2) within each *line*. Contrasts M-P were second-order contrasts conducted between select contrasts; M and N were conducted between-*line*, and O and P within-*line*. CI = 95% Confidence Interval, *conc* = concentration, *temp* = temperature, B6 = C57BL/6J, KO = TRPM8-KO, * = p < 0.05. Note, Contrast Estimate = Condition 1 Estimate - Condition 2 Estimate.

| Contrast ID | Test | Contrast Description | Solution 1 | Solution 1 Estimate [CI] | Solution 2 | Solution 2 Estimate [CI] | Contrast Estimate [CI] | F<sub>(1,1671)</sub> | p-value |
|---|---|---|---|---|---|---|---|---|---|
| A | 1 | Between *temp*<br>within B6 | 100 mM @<br>15°C | 3.37<br>[2.22, 4.91] | 100 mM @<br>30°C | 3.22<br>[2.12, 4.70] | 0.15<br>[-0.58, 0.90] | 0.17 | 0.6842 |
| B | 1 | Between *temp*<br>within KO | 100 mM @<br>15°C | 3.54<br>[2.36, 5.06] | 100 mM @<br>30°C | 2.44<br>[1.57, 3.64] | 1.10<br>[0.43, 1.90] | 11.04 | 0.0009 * |
| C | 2 | Between *temp*<br>within B6 | 500 mM @<br>15°C | 10.24<br>[7.27, 14.27] | 500 mM @<br>30°C | 9.49<br>[6.77, 13.11] | 0.75<br>[-1.04, 2.70] | 0.67 | 0.4401 |
| D | 2 | Between *temp*<br>within KO | 500 mM @<br>15°C | 4.67<br>[3.21, 6.59] | 500 mM @<br>30°C | 3.22<br>[2.14, 4.67] | 1.45<br>[0.62, 2.45] | 12.55 | 0.0006 * |
| E | 3 | Between *conc:temp*<br>within B6 | 100 mM @<br>30°C | 3.27<br>[2.14, 4.80] | 500 mM @<br>15°C | 13.56<br>[9.77, 18.74] | -10.29<br>[-14.23, -7.39] | 210.15 | 9.7x10<sup>-44</sup> * |
| F | 3 | Between *conc:temp*<br>within KO | 100 mM @<br>30°C | 2.49<br>[1.59, 3.68] | 500 mM @<br>15°C | 6.34<br>[4.44, 8.79] | -3.86<br>[-5.43, -2.63] | 80.28 | 2.7x10<sup>-18</sup> * |
| G | 4 | Between *conc:temp*<br>within B6 | 100 mM @<br>15°C | 3.34<br>[2.20, 4.84] | 500 mM @<br>30°C | 12.32<br>[8.86, 17.03] | -8.98<br>[-12.49, -6.39] | 176.16 | 1.1x10<sup>-37</sup> * |
| H | 4 | Between *conc:temp*<br>within KO | 100 mM @<br>15°C | 3.51<br>[2.35, 5.04] | 500 mM @<br>30°C | 4.36<br>[2.98, 6.15] | -0.86<br>[-1.74, -0.05] | 4.39 | 0.0447 * |
| I | 5 | Between *conc*<br>within B6 | 100 mM @<br>15°C | 3.81<br>[2.52, 5.47] | 500 mM @<br>15°C | 14.83<br>[10.59, 20.23] | -11.02<br>[-15.16, -7.64] | 198.46 | 9.0x10<sup>-42</sup> * |
| J | 5 | Between *conc*<br>within KO | 100 mM @<br>15°C | 4.00<br>[2.69, 5.67] | 500 mM @<br>15°C | 6.99<br>[4.84, 9.67] | -2.99<br>[-4.53, -1.77] | 31.79 | 4.6x10<sup>-8</sup> * |
| K | 6 | Between *conc*<br>within B6 | 100 mM @<br>30°C | 3.82<br>[2.59, 5.53] | 500 mM @<br>30°C | 14.33<br>[10.27, 20.03] | -10.51<br>[-14.68, -7.48] | 187.13 | 9.9x10<sup>-40</sup> * |
| L | 6 | Between *conc*<br>within KO | 100 mM @<br>30°C | 2.93<br>[1.90, 4.35] | 500 mM @<br>30°C | 5.17<br>[3.50, 7.34] | -2.24<br>[-3.49, -1.32] | 29.33 | 1.4x10<sup>-7</sup> * |
| M | 3 | Contrasts E, F<br>between *line* | Contrast E | -10.29<br>[-14.23, -7.39] | Contrast F | -3.86<br>[-5.43, -2.63] | -6.43<br>[-10.17, -3.18] | 20.54 | 1.0x10<sup>-5</sup> * |
| N | 4 | Contrasts G, H<br>between *line* | Contrast G | -8.98<br>[-12.49, -6.39] | Contrast H | -0.86<br>[-1.74, -0.05] | -8.13<br>[-11.55, -5.39] | 79.93 | 2.7x10<sup>-18</sup> * |
| O | 3 | Contrasts H, F<br>within KO | Contrast H | -0.86<br>[-1.74, -0.05] | Contrast F | -3.86<br>[-5.43, -2.63] | 3.00<br>[1.67, 4.58] | 23.58 | 2.3x10<sup>-6</sup> * |
| P | 4 | Contrasts G, E<br>within B6 | Contrast G | -8.98<br>[-12.49, -6.39] | Contrast E | -10.29<br>[-14.23, -7.39] | 1.31<br>[-1.28, 4.11] | 0.75 | 0.4401 |

**Figure 5.**
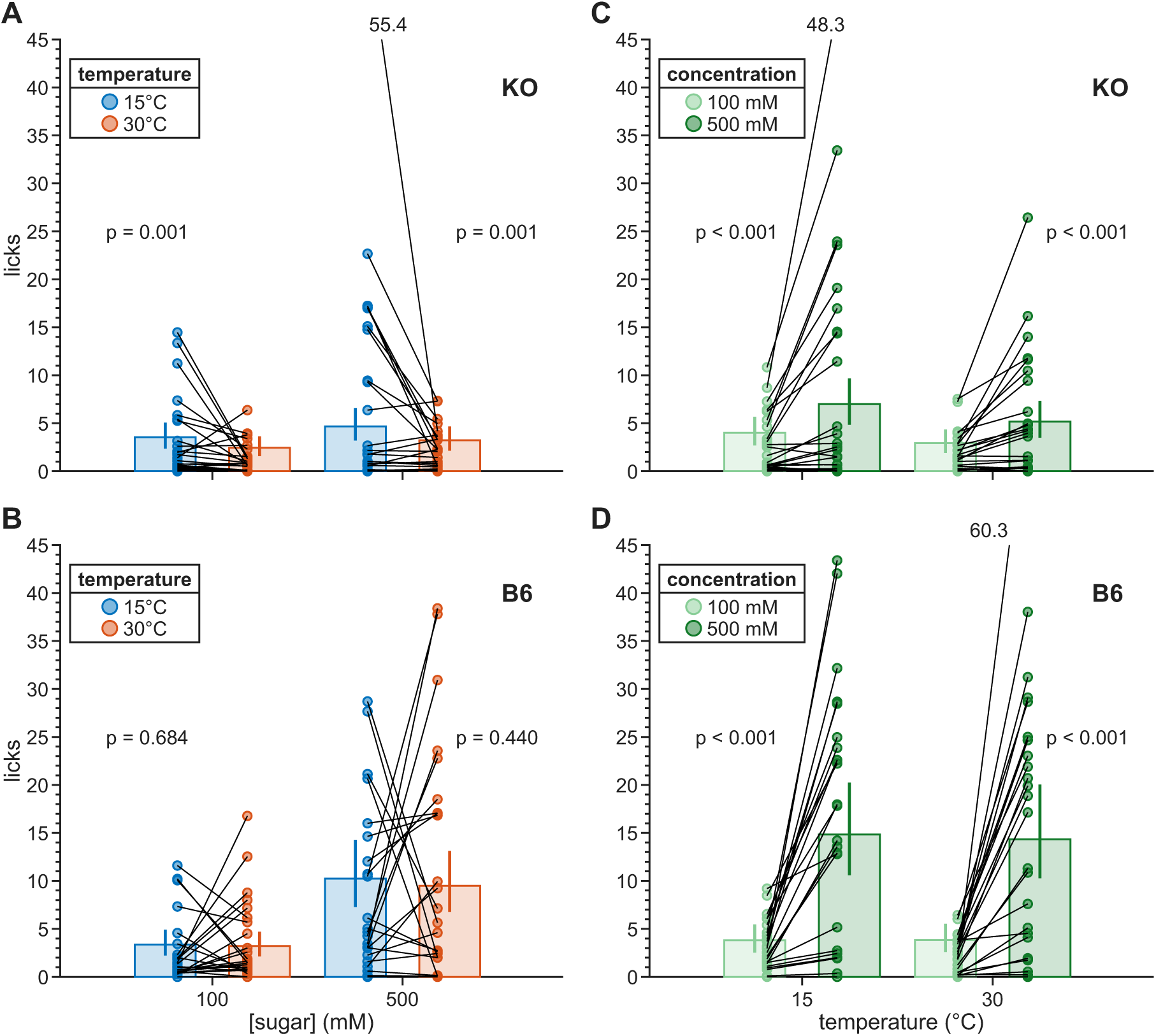
Lick responses within each mouse line to sugar solutions of different temperatures at constant concentrations (A-B) and different concentrations at constant temperatures (C-D). The observed mean licks per solution are plotted as circles, one for each mouse; lines connect observations from the same mouse between the two solutions. The estimated marginal means derived from these data are overlaid as columns. Error bars indicate 95% confidence intervals. (A) Results from Tests 1 and 2 (Table 2), showing responses to a given concentration (100mM or 500mM) when presented at two temperatures (15°C and 30°C) within KO mice. (B) Same as A but for B6 mice. (C) Results from Tests 5 and 6 (Table 2), showing responses to two concentrations when presented at the same temperature for KO mice. (D) Same as C but for B6 mice.

##### 3.2.1.2. Only concentration varies

We observed a significant effect of concentration in both KO and B6 mice (**Table 6 I-L**). At 15°C (**Table 2**, Test 5), B6 mice emitted 11.02 [7.64, 15.16] (F_(1,1671)_ = 198.46, p = 9.0x10^-42^; **Figure 5D**) and KO mice emitted 2.99 [1.77, 4.53] (F_(1,1671)_ = 31.79, p = 4.6x10^-8^; **Figure 5C**) additional licks to 500 mM than 100mM. At 30°C (**Table 2**, Test 6) the contrasts were similar: B6 mice emitted 10.51 [7.48, 14.68] (F_(1,1671)_ = 187.13, p = 9.9x10^-49^; **Figure 5D**) and KO mice emitted 2.24 [1.32, 3.49] (F_(1,1671)_ = 29.33, p = 1.4x10^-7^; **Figure 5C**) additional licks to 500 mM than 100 mM solutions.

##### 3.2.1.3. Both concentration and temperature vary

Next, we asked if varying concentration and temperature inversely (**Table 2**, Test 3) – i.e., low-concentration, higher-temperature (100 mM @ 30°C) presented alongside high-concentration, lower-temperature (500 mM @ 15°C) – evoked similar licks to each sugar solution (**Table 6 E-F**). We observed that both B6 and KO emitted significantly more licks to the ‘high conc - low temp’ solution than the ‘low conc – high temp’ solution: 10.29 [7.39, 14.23] for B6 (*F*_(1,1671)_ = 210.15, *p* = 9.7 × 10^-44^) and 3.86 [2.63, 5.43] for KO (*F*_(1,1671)_ = 80.28, *p* = 2.7 × 10^-18^) (**Figure 6A**).

In addition, we evaluated if lick behavior could be modulated by varying temperature and concentration in parallel (**Table 2**, Test 4) – i.e., ‘low conc - low temp’ (100 mM @ 15°C) presented alongside ‘high conc – high temp’ (500 mM @ 30°C) (**Table 6 G-H**). We observed a similarly significant pattern as the inverse design, with B6 mice emitting 8.98 [6.39, 12.49] and KO mice emitting 0.86 [0.05, 1.74] more licks to the ‘high conc – high temp’ solution (*F*_(1,1671)_ ≥ 4.39 *p* ≤ 0.0447) (**Figure 6B**).

A pair of second order between-line contrasts were conducted within Test 3 and Test 4. In each Test, we contrasted the contrast estimates for each mouse line to determine if the difference in their responses to the two test solutions significantly varied between B6s and KOs. While both lines emitted more licks to the ‘high conc – low temp’ (500 mM @ 15°C) over the ‘low conc – high temp’ (100mM @ 30°C) solutions in Test 3, the between-solution difference was 6.43 [3.18, 10.17] licks larger in B6 mice than KOs (*F*_(1,1671)_ = 20.54, *p* = 1.0 × 10^-5^; **Table 6 M**; **Figure 6A**). Similarly, both lines significantly preferred ‘high conc – high temp’ (500mM @ 30C) solutions over the ‘low conc – low temp’ (100mM @ 15C) in Test 4, but the difference was significantly larger in B6 mice by 8.13 [5.39, 11.55] licks (*F*_(1,1671)_ = 79.93, *p* = 2.7 × 10^-18^; **Table 6 N**; **Figure 6B**).

**Figure 6.**
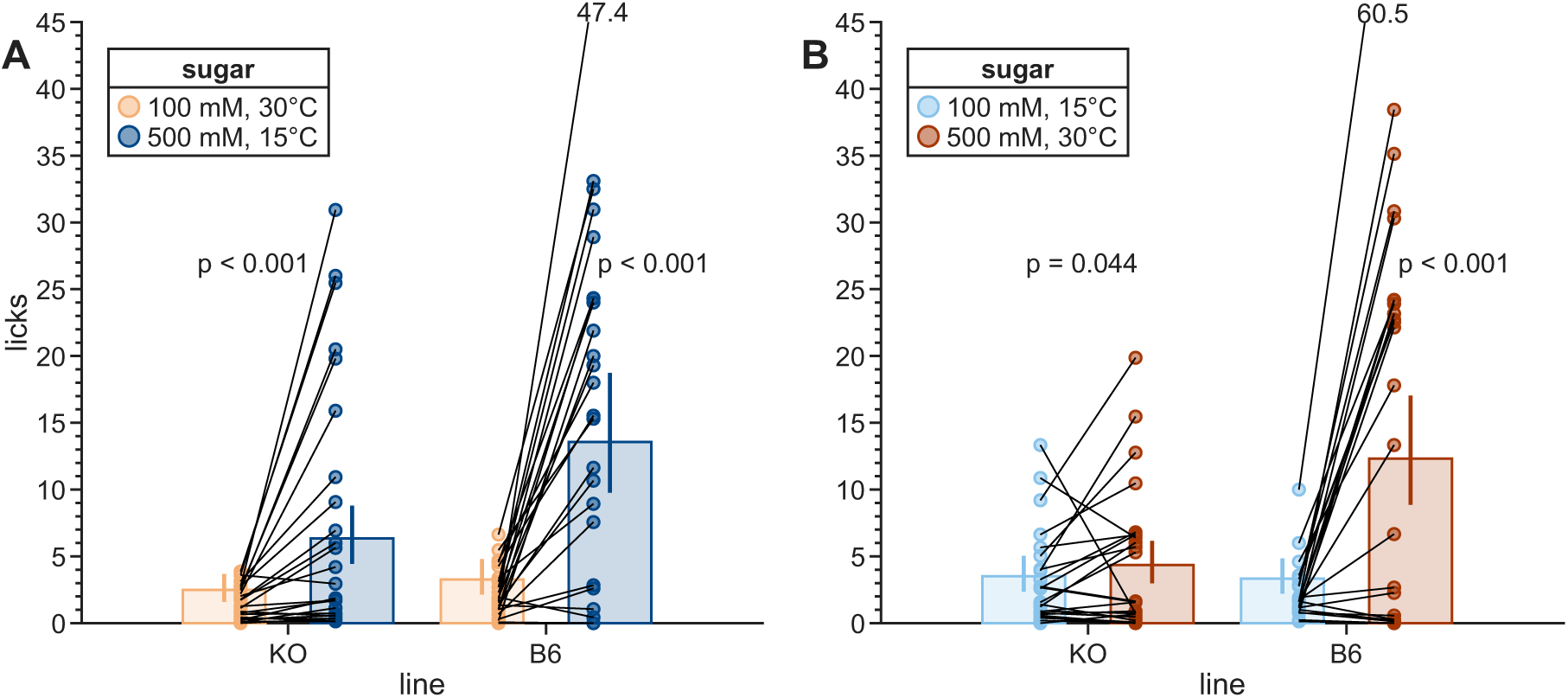
Lick responses within each mouse line to sugar solutions of different temperatures and concentrations. The observed mean licks per solution are plotted as circles, one for each mouse; lines connect observations from the same mouse between the two solutions. The estimated marginal means derived from these data are overlaid as columns. Error bars indicate 95% confidence intervals. (A) Results from Test 3 (Table 2), showing responses to solutions with inversely varied concentrations and temperatures. (B) Results from Test 4 (Table 2), showing responses to solutions with parallel concentrations and temperatures.

Finally, a pair of second order within-line contrasts were conducted to examine if differences in responses to sugar concentrations were the same between the inverse (**Table 2**, Test 3) and parallel (**Table 2**, Test 4) stimulus conditions for each mouse line. B6 mice showed no significant difference in their responses to 100 mM and 500 mM sugars between these Tests (estimate: 1.31 [-1.28, 4.11], *F*_(1,1671)_ = 0.75, *p* = 0.4401; **Table 6 P**, **Figure 6A-B**). In contrast, KO mice had a significantly larger between-solution gap in licking 100 mM and 500 mM sugars when temperature varied inversely with concentration than in parallel (estimate: 3.00 [1.67, 4.58], *F*_(1,1671)_ = 23.58, *p* = 2.3 × 10^-6^, **Table 6 O**, **Figure 6A-B**). In other words, KO mice showed greater avidity to lick 500 mM sugars at 15°C in Test 3 than at 30°C in Test 4.

Altogether, these data suggest that concentration drives licks to sugar more than temperature under the present test conditions. Nevertheless, temperature does have a significant effect that is present only in KO mice, who lick 15°C more than 30°C sugar solutions. In addition, B6 appeared to respond more strongly to a higher sugar concentration than KO mice, both directly and in the magnitude of their preference for a higher concentration solution.

#### 3.2.2. Concentration context inffuences lick behavior to sugar solutions

Next, we asked if mice would emit the same number of licks to the same sugar solution presented in different experimental contexts (i.e., Tests); specifically, when presented alongside sugar solutions of different concentrations and/or temperatures. Our experimental design consisted of four solutions with unique concentration-temperature properties: 100 mM @ 15°C, 100 mM @ 30°C, 500 mM @ 15°C, and 500 mM @ 30°C. Every solution was presented in three Tests, each time paired with one of the other three solutions (**Table 2**). This allowed us to generate EMM licks to a given ‘reference solution’ in each of its three experimental contexts (**Table S6**), which we then contrasted to determine if responses to the reference solution significantly varied based on the context solution it was provided alongside. The specific contrasts referenced below for sugar are detailed in **Table 7** and **Figure 7**. While significant differences in emitted licks under different concentrations were observed in B6 and KO mice, as above, the between-reference solution contrasts did not vary between B6 and KO mice due to the lack of a significant *line:test* interaction in the sugar model, and we did not analyze lines separately.

**Table 7.** Outcomes of contrasts between the estimated marginal mean licks for a given sugar reference solution when paired alongside each of the other three context solutions as indicated by its Test ID (see Table 2). Each row reports the estimated marginal mean [95% confidence interval] licks to a given reference solution when it was presented alongside one of its three possible contexts. Pairwise contrasts were conducted between each of the possible pairings of a given reference solution’s three contexts. CI = 95% Confidence Interval, * = p < 0.05. Note, Contrast Estimate = First Reference Solution Estimate - Second Reference Solution Estimate.

| Contrast ID | Reference Solution | Context Solution | Reference Solution Estimate [CI] | Contrast Estimate [CI] | F<sub>(1,1671)</sub> | p-value | Test ID |
|---|---|---|---|---|---|---|---|
| A | 100 mM @ 15°C | 100 mM @ 30°C | 3.45 [2.66, 4.49] | -0.45 [-1.15, 0.28] | 1.75 | 0.2483 | 1 |
|  |  | 500 mM @ 15°C | 3.90 [2.99, 4.98] |  |  |  | 5 |
| B | 100 mM @ 15°C | 500 mM @ 30°C | 3.42 [2.56, 4.41] | -0.03 [-0.68, 0.56] | 0.01 | 0.9182 | 4 |
|  |  | 100 mM @ 30°C | 3.45 [2.66, 4.49] |  |  |  | 1 |
| C | 100 mM @ 15°C | 500 mM @ 30°C | 3.42 [2.56, 4.41] | -0.48 [-1.19, 0.21] | 2.04 | 0.2309 | 4 |
|  |  | 500 mM @ 15°C | 3.90 [2.99, 4.98] |  |  |  | 5 |
| D | 100 mM @ 30°C | 500 mM @ 30°C | 3.35 [2.50, 4.36] | 0.54 [-0.05, 1.14] | 3.36 | 0.1342 | 6 |
|  |  | 100 mM @ 15°C | 2.81 [2.08, 3.76] |  |  |  | 1 |
| E | 100 mM @ 30°C | 500 mM @ 15°C | 2.86 [2.14, 3.76] | 0.05 [-0.49, 0.59] | 0.03 | 0.9182 | 3 |
|  |  | 100 mM @ 15°C | 2.81 [2.08, 3.76] |  |  |  | 1 |
| F | 100 mM @ 30°C | 500 mM @ 30°C | 3.35 [2.50, 4.36] | 0.49 [-0.14, 1.11] | 2.76 | 0.1659 | 6 |
|  |  | 500 mM @ 15°C | 2.86 [2.14, 3.76] |  |  |  | 3 |
| G | 500 mM @ 15°C | 500 mM @ 30°C | 6.98 [5.48, 8.89] | -3.26 [-4.85, -1.79] | 22.28 | 1.5x10<sup>-5</sup><br>* | 2 |
|  |  | 100 mM @ 15°C | 10.24 [8.06, 12.75] |  |  |  | 5 |
| H | 500 mM @ 15°C | 500 mM @ 30°C | 6.98 [5.48, 8.89] | -2.36 [-3.86, -1.12] | 12.70 | 0.0015<br>* | 2 |
|  |  | 100 mM @ 30°C | 9.34 [7.45, 11.75] |  |  |  | 3 |
| I | 500 mM @ 15°C | 100 mM @ 30°C | 9.34 [7.45, 11.75] | -0.90 [-2.47, 0.73] | 1.33 | 0.2995 | 3 |
|  |  | 100 mM @ 15°C | 10.24 [8.06, 12.75] |  |  |  | 5 |
| J | 500 mM @ 30°C | 100 mM @ 30°C | 8.73 [6.89, 11.06] | 3.07 [1.78, 4.50] | 27.33 | 2.3x10<sup>-6</sup><br>* | 6 |
|  |  | 500 mM @ 15°C | 5.65 [4.43, 7.14] |  |  |  | 2 |
| K | 500 mM @ 30°C | 500 mM @ 15°C | 5.65 [4.43, 7.14] | -1.80 [-2.98, -0.82] | 10.83 | 0.0031<br>* | 2 |
|  |  | 100 mM @ 15°C | 7.45 [5.88, 9.42] |  |  |  | 4 |
| L | 500 mM @ 30°C | 100 mM @ 30°C | 8.73 [6.89, 11.06] | 1.28 [-0.10, 2.76] | 3.75 | 0.1272 | 6 |
|  |  | 100 mM @ 15°C | 7.45 [5.88, 9.42] |  |  |  | 4 |

**Figure 7.**
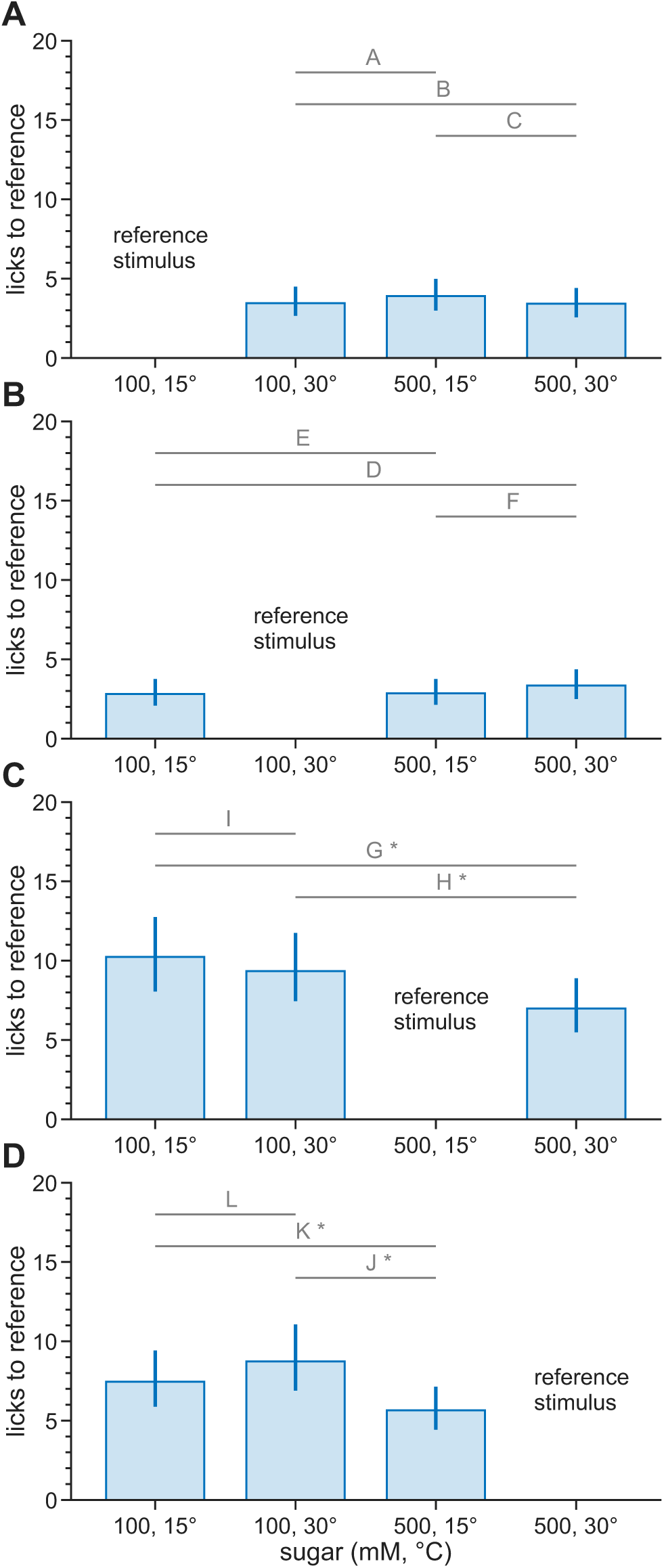
Estimated marginal means of licks to each reference sugar solution in its three contexts. Each subplot (A-D) corresponds to one of the four reference conditions. Bars show licks to the reference solution (y-axes) when presented alongside each of the three context solutions (x-axes). Horizontal lines and letters denote specific contrast tests performed (Table 7). Error bars represent the 95% confidence interval. * = p < 0.05.

Mice emitted 3.42 [2.56, 4.41] licks to **100 mM @ 15**°**C** when presented alongside the higher-concentration higher-temperature 500 mM @ 30°C, 3.90 [2.99, 4.98] licks alongside the higher-concentration 500 mM @ 15°C, and 3.45 [2.66, 4.49] licks alongside the higher-temperature 100 mM @ 30°C. No significant differences were observed between these estimates (*F*_(1,1671)_ ≤ 2.04, *p* ≥ 0.2309; Contrasts A-C; **Figure 7A**).

Mice emitted 2.86 [2.14, 3.76] licks to **100 mM @ 30°C** when presented alongside the higher-concentration lower-temperature 500 mM @ 15°C, 3.35 [2.50, 4.36] licks alongside the higher-concentration 500 mM @ 30°C, and 2.81 [2.08, 3.76] licks alongside lower-temperature 100 mM @ 15°C. No significant differences were observed between these estimates (*F*_(1,1671)_ ≤ 3.36, *p* ≥ 0.1342; Contrasts D-F; **Figure 7B**).

Mice emitted 6.98 [5.48, 8.89] licks to **500 mM @ 15°C** when presented alongside the higher-temperature 500 mM @ 30°C, 9.34 [7.45, 11.75] licks alongside the lower-concentration higher-temperature 100 mM @ 30°C, and 10.24 [8.06, 12.75] licks when tested with the lower-concentration 100 mM @ 15°C. Mice emitted significantly fewer licks to **500 mM @ 15°C** when it was presented alongside 500 mM @ 30°C than when presented alongside 100 mM @ 15°C (*F*_(1,1671)_ = 22.28, *p* = 1.5 × 10^-5^; Contrast G) or 100 mM @ 30°C (*F*_(1,1671)_ = 12.70, *p* = 0.0015; Contrast H), reflecting a concentration effect. No significant difference was observed between the two 100 mM contexts (*F*_(1,1671)_ = 1.33, *p* = 0.2995; Contrast I; **Figure 7C**).

Mice emitted 5.65 [4.43, 7.14] licks to **500 mM @ 30°C** when presented alongside the lower-temperature 500 mM @ 15°C, 8.73 [6.89, 11.06] licks alongside the lower- concentration 100 mM @ 30°C, and 7.45 [5.88, 9.42] licks alongside the lower-concentration lower-temperature 100 mM @ 15°C. Mice made significantly fewer licks to **500 mM @ 30°C** when it was presented alongside 500 mM @ 15°C than when presented alongside 100 mM @ 15°C (*F*_(1,1671)_ = 10.83, *p* = 0.0031; Contrast K) or 100 mM @ 30°C (*F*_(1,1671)_ = 27.33, *p* = 2.3 × 10^-6^; Contrast J). No significant difference was observed between the two 100 mM contexts (*F*_(1,1671)_ = 3.75, *p* = 0.1272; Contrast L; **Figure 7D**).

Altogether, our results indicate that mice emitted significantly more licks to 500 mM when the alternative was a 100 mM solution, regardless of temperature. In contrast, responses to 100 mM solutions did not significantly vary between different contexts, even when presented alongside 500 mM solutions. When a 100 mM and 500 mM solution were presented together, licks emitted to 100 mM did not change, but licks to 500 mM increased. In other words, the observed context-dependent changes in licks appear to be driven by a preference for higher sugar concentrations.

**Table 8.** In order to evaluate line- and concentration-dependent changes in licks to sugars across experimental exposures, a total of eight simple effect contrasts were conducted between the estimated marginal means for each of the four line:concentration groups on the first (0) and last (15) experimental exposures (*exp*). For each contrast, the estimated marginal means, their estimated contrast, the estimated 95% confidence intervals, the F-statistic, and the FDR-corrected p-value are reported. All contrasts = Condition 1 Estimate - Condition 2 Estimate. CI = 95% Confidence Interval, *exp* = exposure, * = p < 0.05.

| Contrast ID | Contrast Description | Condition 1 | Condition 1 Estimate [CI] | Condition 2 | Condition 2 Estimate [CI] | Contrast Estimate [CI] | F<sub>(1,1671)</sub> | p-value |
|---|---|---|---|---|---|---|---|---|
| A | B6, 500 mM v 100 mM, *exp* 0 | B6 500 mM 0 | 8.99 [6.37, 12.72] | B6 100 mM 0 | 3.26 [2.15, 4.81] | 5.73 [3.91, 8.59] | 66.47 | 5.1x10<sup>-15</sup> * |
| B | B6, 500 mM v 100 mM, *exp* 15 | B6 500 mM 15 | 16.62 [12.07, 22.81] | B6 100 mM 15 | 3.92 [2.70, 5.78] | 12.70 [9.02, 17.69] | 218.36 | 1.7x10<sup>-45</sup> * |
| C | KO, 500 mM v 100 mM, *exp* 0 | KO 500 mM 0 | 5.16 [3.58, 7.36] | KO 100 mM 0 | 3.06 [1.99, 4.51] | 2.10 [0.97, 3.47] | 16.31 | 0.0001 * |
| D | KO, 500 mM v 100 mM, *exp* 15 | KO 500 mM 15 | 5.25 [3.60, 7.46] | KO 100 mM 15 | 3.42 [2.32, 5.01] | 1.83 [0.90, 2.96] | 16.65 | 9.4x10<sup>-5</sup> * |
| E | KO v B6, 100 mM, *exp* 0 | KO 100 mM 0 | 3.06 [1.99, 4.51] | B6 100 mM 0 | 3.26 [2.15, 4.81] | -0.20 [-2.03, 1.64] | 0.05 | 0.8610 |
| F | KO v B6, 100 mM, *exp* 15 | KO 100 mM 15 | 3.42 [2.32, 5.01] | B6 100 mM 15 | 3.92 [2.70, 5.78] | -0.50 [-2.65, 1.36] | 0.26 | 0.7039 |
| G | KO v B6, 500 mM, *exp* 0 | KO 500 mM 0 | 5.16 [3.58, 7.36] | B6 500 mM 0 | 8.99 [6.37, 12.72] | -3.83 [-8.03, -0.39] | 4.89 | 0.0459 * |
| H | KO v B6, 500 mM, *exp* 15 | KO 500 mM 15 | 5.25 [3.60, 7.46] | B6 500 mM 15 | 16.62 [12.07, 22.81] | -11.37 [-17.47, -6.34] | 24.21 | 2.6x10<sup>-6</sup> * |

#### 3.2.3. Licks to sugars increase with experimental exposures at high concentrations in a line-dependent manner

During model fitting, we observed a significant longitudinal effect on licking behavior across the study period, whereby estimated licks increased with every additional day of testing (i.e., ‘exposure’) in a *line*- and concentration-(*conc*) dependent manner (**Table S4**; **Equation 2**). We explored this by quantifying the longitudinal effect of exposure (*exp*) within each of the four unique *line*:*conc* groups (B6:100 mM, B6:500 mM, KO:100 mM, KO:500 mM) using EMMs, and derived contrast estimates between each of these groups to evaluate their relative magnitude and significance.

Due to the structural interdependence of *exp* and *blockDay* (**Table S4**; **Equation 2**), we generated EMMs only for the first *exp* to each Test (i.e. every 3rd *exp*, where *blockDay* = 0, starting at *exp 0*), for a total of six estimates per group on *exps* 0, 3, 6, 9, 12, and 15. The EMMs and their associated contrasts for the first (0) and last (15) *exp* are reported in **Table 8**. The estimated slopes within each group are reported in **Table 9** and plotted in **Figure 8**.

**Figure 8.**
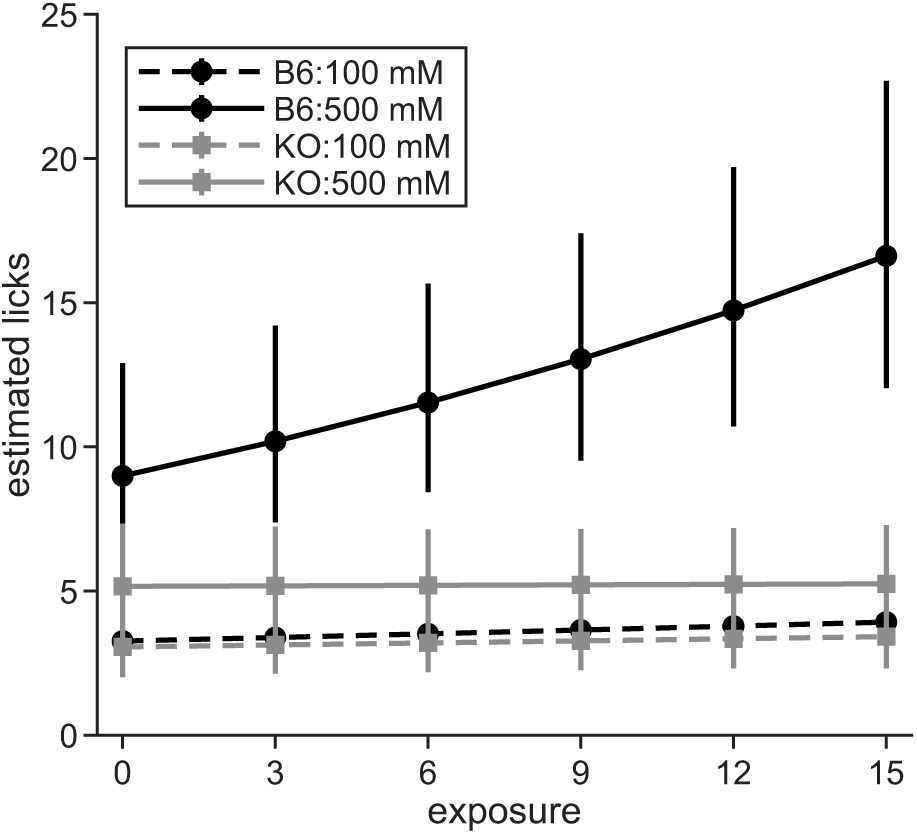
Estimated marginal mean licks for each exposure, plotted by line:concentration. Error bars represent 95% confidence intervals. Estimated slopes were derived via the average rate of change between estimates on exp 0 and *exp* 15.

Both B6 and KO mice emitted significantly more licks to 500 mM than 100 mM sugars throughout the course of the study (**Table 8**). B6 mice made 5.73 [3.91, 8.59] (*F*_(1,1671)_ = 66.47, *p* = 5.1 × 10^-15^) more licks to 500 mM than 100 mM on *exp* 0, which increased to 12.70 [9.02, 17.69] (*F*_(1,1671)_ = 218.36, *p* = 1.7 × 10^-45^) on *exp* 15 (**Table 8 A-B**). Similarly, KO mice made 2.10 [0.97, 3.47] (*F*_(1,1671)_ = 16.31, *p* = 0.0001) more licks to 500 mM than 100 mM on *exp* 0 (**Table 8 C-D**). Yet while the contrast remained significant on *exp* 15 (*F*_(1,1671)_ = 16.65, *p* = 9.4 × 10^-5^), the magnitude of the contrast decreased to 1.83 [0.90, 2.96].

**Table 9.** To estimate the rate of change in licks to sugars per experimental exposure (*exp*), we conducted four simple effect contrasts, one for each line:concentration group, between the last (15) and first (0) *exp*. The resulting contrast estimates and CIs were then divided by 15 to convert them into a ‘licks per exposure’ slope estimate. For each contrast, the estimated marginal means, the adjusted estimated contrast (i.e., slope), the estimated 95% confidence intervals, the F-statistic, and the FDR-corrected p-value are reported. CI = 95% Confidence Interval, *exp* = Exposure, * = p < 0.05.

| Contrast Description | *exp* 15 Estimate [CI] | *exp* 0 Estimate [CI] | Slope Estimate [CI] | F<sub>(1,1671)</sub> | p-value |
|---|---|---|---|---|---|
| B6 100 mM, Slope | 3.920 [2.701, 5.776] | 3.262 [2.153, 4.813] | 0.044 [-0.020, 0.120] | 1.76 | 0.2535 |
| B6 500 mM, Slope | 16.619 [12.066, 22.811] | 8.990 [6.372, 12.716] | 0.509 [0.285, 0.800] | 25.31 | 1.7x10<sup>-6</sup> * |
| KO 100 mM, Slope | 3.417 [2.324, 5.012] | 3.058 [1.991, 4.507] | 0.024 [-0.042, 0.093] | 0.59 | 0.5413 |
| KO 500 mM, Slope | 5.247 [3.596, 7.457] | 5.160 [3.580, 7.363] | 0.006 [-0.082, 0.096] | 0.02 | 0.8970 |

The difference between B6 and KO mice varied by concentration (**Table 8 E-H**). At 100 mM, no significant difference was observed between the lines: B6s emitted 0.20 [-1.64, 2.03] (*F*_(1,1671)_ = 0.05, *p* = 0.8610) more licks than KO mice on *exp* 0, and 0.50 [-1.36, 2.65] (*F*_(1,1671)_ = 0.26, *p* = 0.7039) more licks on *exp* 15. At 500 mM, we observed significant between-line differences. B6s emitted 3.83 [0.39, 8.03] (*F*_(1,1671)_ = 4.89, *p* = 0.0459) more licks than KOs on *exp* 0, and 11.37 [6.34, 17.47] (*F*_(1,1671)_ = 24.21, *p* = 2.6 × 10^-6^) more licks than KOs on *exp* 15.

The largest change in magnitude over the course of the study (*exp* 0 to *exp* 15) was observed in B6 mice at 500 mM (+7.63; **Table 9**). The other groups experienced comparatively smaller changes: B6 mice at 100 mM increased by 0.66 licks, KO mice at 100 mM increased by 0.36 licks, and KO mice at 500 mM increased by 0.09 licks. These results support our previous observations that B6 mice tend to respond more strongly to higher concentrations than KO mice (**Table 6**) and suggest this is dependent upon *exp*.

We estimated the slopes of the effect of *exp* in each *line:conc* group (**Table 9**). A significant effect of *exp* was observed in B6 mice at 500 mM, whereby each *exp* was associated with 0.509 [0.285, 0.800] (*F*_(1,1671)_ = 25.31, *p* = 1.7 × 10^-6^) additional licks (**Table 9**). However, this effect became non-significant at the 100 mM concentration, with only 0.044 [-0.020, 0.120] (*F*_(1,1671)_ = 1.76, *p* = 0.2535) additional licks per *exp*. KOs had non-significant slopes at both concentrations: at 100 mM, licks increased by 0.024 [-0.042, 0.093] (*F*_(1,1671)_ = 0.59, *p* = 0.5413) per *exp*, and at 500 mM the slope became 0.006 [-0.082, 0.096] (*F*_(1,1671)_ = 0.02, *p* = 0.8970) licks per *exp*.

Altogether these data indicate that B6 mice, but not KOs, increase their licks to sugar across repeated exposures. This effect was only observed at the 500 mM concentration, which may suggest that the effect scales with concentration, or only manifests once a certain concentration threshold is reached.

## 4. Discussion

### 4.1. Temperature and quinine acceptance

Our analysis indicates that various factors influence licking activity to quinine. Concentration and temperature both modulate responses to quinine solutions in an apparently independent yet summative fashion, with higher concentration (0.3 mM) and/or higher temperature (30°C) being associated with fewer licks to (i.e., less acceptance of) quinine. In addition, acceptance is further modulated by experimental context – mice generally made the smallest number of licks to a quinine solution when the alternative was a lower temperature (15°C) and/or concentration (0.03 mM). Finally, quinine acceptance increases longitudinally with each subsequent exposure to experimental activities. This may reflect multiple effects, such as increasing thirst pressure due to water restriction, reduced stress due to habituation to the experimental protocol, or a learned association between experimental procedures and thirst relief. Moreover, all these effects appear to be independent of TRPM8 genotype, as a significant mouse line effect was not found.

Our results were observed using two fluid temperatures, 30°C and 15°C, that rely on TRPM8 for oral sensation but stimulate equal licks (indifference) in thirst-motivated B6 mice when paired with water in brief-access tests (Li et al., 2024). Brief-access data from water-restricted rats show similar indifference to lick water at 30°C and lower temperatures (e.g., 10°C; Torregrossa et al., 2012). That the present addition of quinine caused 30°C to become avoided in B6 mice agrees with the proximity of 30°C to higher temperatures that reduce water licking in this line in brief-access tests (35°C to 36°C; Li et al., 2024; Marques et al., 2026), which may have allowed quinine to cause a stronger reduction in the palatability of the 30°C than 15°C solution.

Evidence for additivity between 30°C and quinine stems from our analysis of experimental context, which identified that concentration and temperature individually affect licks to a given quinine solution. Specifically, presenting a reference solution alongside one with a higher concentration or a higher temperature will increase licks to the reference solution, but to a lesser extent than when presented against a condition where both concentration and temperature are higher. There was a stronger effect of temperature on quinine licking, as evinced by larger contrast estimates for the temperature-varying (**Table 4**, Contrasts C, F, I, and L) versus concentration-varying (**Table 4**, Contrasts B, E, H, and K) conditions. What is more, the largest contrast estimates in the contextual analysis were observed when concentration and temperature were varied in parallel (i.e., both increased or decreased; **Table 4**, Contrasts D and G), revealing a compounding effect of temperature and concentration on quinine choice.

These observations highlight a complexity to capturing temperature effects on taste preference behavior where available temperature and concentration options, and test parameters, can impact outcomes. Historically, brief-access tests have monitored licking across several room-temperature concentrations of a taste chemical presented in one test session along with water. The present studies purposefully omitted water from quinine testing to force mice to choose between only two bitter fluids with differing physical properties when faced with thirst pressure, which interacts with taste solution valence to affect licking (Spector & St John, 1998). Speculatively, consumption tests that proffer only adulterated fluids may have relevance to ecological settings where animals must consume less palatable and variably toxic substances to maintain homeostasis (Glendinning et al., 2006; Rowland & Flamm, 1977; Spector & St John, 1998). Including water presentations within our quinine tests may yield different results (Glendinning, 1993) and complicate interpretations of some conditions (e.g., inverse or parallel concentration and temperature).

That mice showed less acceptance of 30°C than 15°C (cooler) quinine accords with human psychophysical data on thermal influences on quinine taste perception. Green and Andrew (2017) found that across a broad range of fluid temperatures (e.g., from 10° to 40°C), cooling suppressed the perceived bitterness of suprathreshold concentrations of quinine relative to warm temperatures. Other data derived from studies that gauged time-dependent dynamics in human taste perception suggest that 35°C quinine may be perceived as more bitter than cold (5°C) during the first few seconds of gustatory stimulation (Bajec et al., 2012). These results suggest that temperature modulates the intensity of quinine’s taste. It is important to acknowledge that the present behavioral analysis can only capture how mice respond to taste stimuli proffered at controlled concentrations and temperatures and cannot address how these parameters effect perceived taste intensity. Nevertheless, neurophysiological studies in rodents show cooling and warming taste solutions can, respectively, decrease and increase electrophysiological activity to quinine in subsets of peripheral (Breza et al., 2006) and central (Wilson & Lemon, 2013) gustatory neurons, implying temperature impacts the intensity of quinine responses in neural circuits for taste.

We have detected a potential confounding effect created by the repeated testing of a mouse within this experimental design (**Figure 4**), where licking acceptance of quinine increased with exposure. This may be due to several individual effects. The mice were water-deprived, therefore each passing day likely increased thirst pressure (St John et al., 2017). The mice may have also habituated to the experimental procedure and apparatus (Staszko et al., 2022; Svedberg et al., 2026), potentially reducing stress as the study progressed. Additionally, the mice may have developed an association between experimental procedures and thirst relief. It is reasonable to infer that each of these effects (higher thirst, reduced stress, learned association) may increase motivation to lick, resulting in the observed exposure effect.

What is more, we found that this exposure effect is dependent upon temperature. Specifically, it is stronger at 15°C than 30°C (**Table 5**). Changes in stimulus temperature may affect the theorized individual effects that summate to the exposure effect. For instance, lower fluid temperatures may induce less stress (Li et al., 2024) and satisfy thirst faster (Torregrossa et al., 2012). Cool water and tongue cooling are discussed to be more satiating than warm (Eccles et al., 2013; Kapatos & Gold, 1972; Mendelson & Chillag, 1970), albeit it is difficult to assert that this concept is reflected in the present results. Behavioral correlates of cold satiation in rodent fluid consumption tests are found more so in long-term, but not brief-access, assays (Kay et al., 2020) and follow a reduction in licking to cool compared to warm fluid temperatures (Kapatos & Gold, 1972; Kay et al., 2020). Some evidence for cold satiation in rodents is based on temperature ranges broader than tested here (Kay et al., 2020; Marques et al., 2026).

While this exposure effect is likely very dependent upon study design (i.e., tastant, solution properties, duration, etc.), this analysis provides an estimate of its magnitude, which can be used to inform the design and analysis of future studies. Importantly, these results provide quantitative evidence of the existence of an effect of repeated testing and prolonged water deprivation on lick rate to quinine and demonstrate that it is worth considering in the analysis of similar studies. The present experiments included a control for repeated testing by randomizing the quinine, and sugar, test order across mice, which resulted in different test conditions/stimulus pairings being tested on different mice each day.

### 4.2. TRPM8 and sugar preference

Unlike quinine, lick rates to sugar solutions were captured from mice that were water replete and not thirst-or hunger-motivated during testing. This was intended to isolate their preference for sugars due to stimulus hedonics, with the brief-access nature of the fluid licking tests capturing orosensory reactions to this stimulus feature. We found that temperature produced a small, but significant, effect on licking to sugars in KO mice. When only temperature varied, KO mice made significantly more licks to 15°C compared to 30°C sugar whereas B6 mice were indifferent. Furthermore, KO mice made significantly more licks to the high (500 mM) than low (100 mM) sugar concentration when temperature varied inversely with concentration (i.e., the high concentration was 15°C and the low was 30°C) opposed to in parallel (i.e., the high was 30°C, and the low 15°C) (**Table 6**, **Figure 6A-B**). These results show higher fluid temperatures reduce licking to sugar in KO mice.

The reduced response to 30°C sugar agrees with our prior data that showed TRPM8 deficient mice lick more 15°C than 30°C water in two-fluid thermolickometry tests, whereas B6 mice are indifferent to these water temperatures (Li et al., 2024). This behavioral difference aligns with analyses of trigeminal neural responses in TRPM8 deficient mice that found an anomalous positive correlation between activity to oral stimulation with temperatures near 30°C and aversively warmer/hot temperatures in TRPM8 deficient mice (Li et al., 2024). We hypothesized that 30°C adopts a negative valence without TRPM8 (Li et al., 2024), with this changed valence presently appearing to compete with sugar taste preferences in KO mice. This concept implies that active TRPM8 channels support the development of oral thermal sensing in ways that interact with taste preferences for sugars and their gustatory appeal. Because TRPM8 is expressed by trigeminal fibers that surround, but do not innervate, taste buds (Abe et al., 2005; Dhaka et al., 2008), orosensory interactions between TRPM8 input and sweet taste messages appear likely to arise downstream of receptor events.

The significant effect of temperature observed in KO mice arose amidst notable variance in responding to sugar across animals. For instance, when only temperature was varied, a subset of KO mice made substantially more licks to 15°C sugars compared to 30°C, with others showing low activity and indifference to respond to either temperature (e.g., **Figure 5A**). This variance may be contributed by a lack of thirst pressure on licking in water-replete mice and/or the *ad libitum* availability of food in their home cage, which may have caused varied consummatory motivational states during testing. Moreover, we observed a tendency for mainly females to show low responsiveness in some sugar tests, albeit a significant effect of sex did not emerge from the omnibus analysis performed on all data (**Table S4**). Finally, mixed behavioral patterns were also observed in B6 mice. Some notably licked more 30°C than 15°C sugar when only temperature varied while others showed the converse or indifference (**Figure 5B**), resulting in a non-significant difference in responding between temperatures. This counters neurophysiological results showing higher gustatory firing to sugars at warmer compared to cooler temperatures in inbred mice (Li & Lemon, 2015; Lu et al., 2016; Lu et al., 2012; Talavera et al., 2005; Wilson & Lemon, 2013, 2014) – data that suggest higher temperatures can increase sugar taste intensity. While some of the present behavioral variance may have stemmed from test conditions, it is curious if species was a factor. Brief-access thermal licking studies in water-replete rats have found a significant effect of temperature on preference for sucrose under certain conditions (Torregrossa et al., 2012).

B6 and KO mice emitted significantly more licks to the high compared to low concentration of sugars when both were offered in one test. Yet, compared to B6, KO mice displayed a smaller increase in licking high sugar concentrations in these settings than was attributable to concentration. This was particularly evident when sugar concentration was varied under constant fluid temperature (**Table 6**, **Figure 5C-D**). What is more, B6 mice longitudinally increased their licking preference for 500 mM sugars with additional exposures whereas KO mice showed no change in responding to sugars over time (**Tables 8**, **9**; **Figure 8**). Thus, KO mice show a generally reduced avidity to lick sugar solutions.

TRPM8 deficiency is established to affect energy balance and feeding under certain conditions. For instance, TRPM8 deficient mice show hyperphagia and increased stored fat mass when housed at ambient temperatures around 21° to 25°C (Reimundez et al., 2018), which fall below thermoneutral temperatures indexed for mice (Gordon, 1993). These effects were accompanied by metabolic changes including increased levels of circulating leptin – an anorexigenic mediator (Reimundez et al., 2018). There is some evidence that increased leptin can suppress taste receptor cell and gustatory nerve responses to sweet substances and decrease licking towards these stimuli in mice (Kawai et al., 2000; Shigemura et al., 2004; Yoshida et al., 2013). Yet leptin effects on mouse gustatory neural and behavioral responses to sweet taste stimuli have not been universally found (Glendinning et al., 2015). Nevertheless, it is curious if some facet of metabolic change in TRPM8 deficient mice is regulating their lowered taste preference for sugars. Additionally, that neural and behavioral responses to ∼30°C anomalously resemble warmer temperatures in TRPM8 deficient mice (Li et al., 2024) may suggest they experience a warmer than normal world, with warm environments capable of suppressing energy intake in mice under select experimental conditions (Kaiser et al., 2026; Morris et al., 2020). As a next step, it would be informative to investigate if TRPM8 deficiency impacts gustatory neurophysiological responses to sucrose and glucose.

These findings contribute to a growing literature on interactions between somatosensation and gustation. Studying taste and thermosensation together appears critical for understanding the operational principles of these senses individually and how they may intersect to drive preference and intake behavior. While prior and the present data indicate that taste sensation is reliant on temperature and thermosensation, the evolutionary and functional bases of this link remain speculative at best. Moreover, the parabrachial nucleus is evidenced to support the integration of aversive taste information with nociceptive thermosensory signals in mice (Li et al., 2022; Li & Lemon, 2019, 2026), albeit a neural mechanism that could mediate interplay between appetitive sugar tastes and thermal messages mediated by TRPM8 remains unknown. Further investigations in this realm will be critical to delineate how diverse sensory factors affect food preference and nutritional status, and for understanding gustatory and somatosensory coding.

## Acknowledgements

Research reported in this publication was supported by NIH grant DC020843 to C.H.L. The content is solely the responsibility of the authors and does not necessarily represent the official views of the NIH. Portions of these data were presented in abstract form at the 2025 meeting of the Association for Chemoreception Sciences. Mehrnoush Nourbakhsh aided with data collection. The authors thank Rosalie Maltby for critical feedback on an earlier draft of this manuscript.

## Data Availability Statement

The data used in statistical analysis and their data dictionary are available through an OSF data repository via the following link: https://osf.io/xw6my/overview?view_only=20916f30932b41a4aed57654b205d6a5

## Author contributions

**Kyle T. Zumpano:** Conceptualization, investigation, methodology, project administration, validation, visualization, writing – original draft, writing – review & editing. **Chris Brooks:** Data curation, formal analysis, methodology, software, validation, visualization, writing – original draft, writing – review & editing. **Christian H. Lemon**: Conceptualization, data curation, formal analysis, funding acquisition, methodology, project administration, resources, software, supervision, validation, visualization, writing – original draft, writing – review & editing.

