## Supplemental tables for "Taste preferences are influenced by temperature and the TRPM8 thermoreceptor"

### Supplemental Materials

| Model Specification | | Effects | | Model Fit Parameters | | | | Likelihood-Ratio Test Outcomes | | | |
|---|---|---|---|---|---|---|---|---|---|---|---|
| Description | ID | Fixed Effects | Random Effects | AIC | BIC | LL | Model DF | Reference Model | Delta DF | $X^2$ | p-value |
| Intercept Only | M0I | N/A | Random Intercept at Mouse level | 2406.3 | 2417.0 | -1200.1 | 3 | N/A | N/A | N/A | N/A |
| Structural | M0 | M0I + *test* + *sex* + *exp* | | 2295.8 | 2331.5 | -1137.9 | 10 | M0I | 7 | 124.45 | < 0.0001 * |
| Core Scientific | M1 | M0 + *line* + *conc* + *temp* | | 2185.5 | 2231.9 | -1079.8 | 13 | M0 | 10 | 240.72 | < 0.0001 * |
| *line:sex* interaction | M2a | M1 + *line:sex* | | 2187.5 | 2237.5 | -1079.8 | 14 | M1 | 1 | < 0.01 | 0.9658 |
| *line:temp* interaction | M2b | M1 + *line:temp* | | 2187.5 | 2237.4 | -1079.7 | 14 | M1 | 1 | 0.09 | 0.7588 |
| *line:conc* interaction | M2c | M1 + *line:conc* | | 2187.4 | 2237.4 | -1079.7 | 14 | M1 | 1 | 0.13 | 0.7235 |
| *sex:temp* interaction | M2d | M1 + *sex:temp* | | 2168.8 | 2236.8 | -1079.4 | 14 | M1 | 1 | 0.74 | 0.3903 |
| *sex:conc* interaction | M2e | M1 + *sex:conc* | | 2187.5 | 2237.4 | -1079.7 | 14 | M1 | 1 | 0.09 | 0.7593 |
| *conc:temp* interaction | M2f | M1 + *conc:temp* | | 2187.1 | 2237.1 | -1079.5 | 14 | M1 | 1 | 0.45 | 0.5020 |
| *line:sex:temp* interactions | M2g | M1 + *line:sex:temp* | | 2192.7 | 2253.4 | -1079.4 | 17 | M1 | 4 | 0.85 | 0.9321 |
| *line:sex:conc* interactions | M2h | M1 + *line:sex:conc* | | 2192.7 | 2253.4 | -1079.4 | 17 | M1 | 4 | 0.83 | 0.9349 |
| *line:conc:temp* interactions | M2i | M1 + *line:conc:temp* | | 2191.9 | 2252.6 | -1079.0 | 17 | M1 | 4 | 1.62 | 0.8044 |
| *sex:conc:temp* interactions | M2j | M1 + *sex:conc:temp* | | 2191.8 | 2252.4 | -1078.9 | 17 | M1 | 4 | 1.78 | 0.7755 |
| *line:conc:sex:temp* interactions | M2k | M1 + *line:conc:sex:temp* | | 2204.1 | 2289.7 | -1078.0 | 24 | M1 | 11 | 3.48 | 0.9827 |
| *test:line* interaction | M2l | M1 + *test:line* | | 2191.6 | 2255.8 | -1077.8 | 18 | M1 | 5 | 3.95 | 0.5572 |

| | | | | | | | | | | | |
|---|---|---|---|---|---|---|---|---|---|---|---|
| *test:sex* interaction | M2m | M1 + *test:sex* | | 2192.4 | 2256.7 | -1078.2 | 18 | M1 | 5 | 3.10 | 0.6839 |
| *line:exp* interaction | M2n | M1 + *line:exp* | | 2185.2 | 2235.1 | -1078.6 | 14 | M1 | 1 | 2.37 | 0.1241 |
| ***temp:exp* interaction** | **M2o** | **M1 + *temp:exp*** | | **2180.5** | **2230.5** | **-1076.3** | **14** | **M1** | **1** | **7.00** | **0.0081 *** |
| *conc:exp* interaction | M2p | M1 + *conc:exp* | | 2187.5 | 2237.5 | -1079.7 | 14 | M1 | 1 | 0.05 | 0.8202 |
| *sex:exp* interaction | M2q | M1 + *sex:exp* | | 2184.6 | 2234.6 | -1078.3 | 14 | M1 | 1 | 2.92 | 0.0873 |

**Table S1** - Model fit parameters and outcomes of likelihood-ratio tests used in model fitting and final model selection for the quinine data. The final analytical model (M2o) is bolded. Total observations = 262; Total subjects = 22; AIC: Aikake Information Criterion; BIC: Bayesian Information Criterion; DF: Degrees of Freedom; LL: Log-Likelihood; N/A: Not Applicable. * = $p < 0.05$

| Model Term | Reference Level | Predictor Level | Estimate | SE | p-value |
|---|---|---|---|---|---|
| $\beta_0$ | All terms at Reference Level (*exp* = 0) | Intercept (Baseline Estimate) | -46.84 | 6.23 | $1.2 \times 10^{-9}$ * |
| $\beta_{test2}$ | Reference Test (Table1: Test 1) | Table 1: Test 2 | 9.10 | 3.43 | 0.0086 * |
| $\beta_{test3}$ | | Table1: Test 3 | -2.03 | 3.17 | 0.5218 |
| $\beta_{test4}$ | | Table1: Test 4 | 11.51 | 3.13 | 0.0003 * |
| $\beta_{test5}$ | | Table1: Test 5 | 3.79 | 2.97 | 0.2033 |
| $\beta_{test6}$ | | Table1: Test 6 | 3.50 | 2.97 | 0.2394 |
| $\beta_{line}$ | B6 | KO | 17.00 | 6.41 | 0.0086 * |
| $\beta_{sex}$ | Female | Male | 1.98 | 6.42 | 0.7583 |
| $\beta_{conc}$ | 0.03 mM | 0.3 mM | -14.92 | 1.99 | $1.1 \times 10^{-12}$ * |
| $\beta_{temp}$ | 15°C | 30°C | -9.24 | 4.02 | 0.0223 * |
| $\beta_{exp}$ | $1^{st}$ Exposure (*exp* = 0) | $2^{nd}$ Exposure (*exp* = 1) | 8.93 | 0.69 | $8.4 \times 10^{-30}$ * |
| $\beta_{temp:exp}$ | 15°C, Exposure 0 | 30°C, Exposure 1 | -2.63 | 0.99 | 0.0082 * |

**Table S2** - Fixed effect coefficient estimates from the final analytical quinine linear mixed model. Estimates, and their standard errors and p-values, are shown for the reference-coded model terms predicting water-adjusted lick responses. Reference Level indicates the reference group for that term. The Estimate represents the estimated change in water-adjusted licks between a Predictor and its Reference Level (or 0, for Intercept), and the p-value the significance of that effect. Intercept represents the estimated response at all Reference Levels and exposure = 0 (i.e. Day 1). SE: Standard Error, * = $p < 0.05$

| Model Specifications | | Effects | | Model Fit Parameters | | | | Likelihood-Ratio Test Outcomes | | | |
|---|---|---|---|---|---|---|---|---|---|---|---|
| Description | ID | Fixed Effects | Random Effects | AIC | BIC | LL | Model DF | Reference Model(s) | Delta DF | $X^2$ | p-value |
| Intercept Only | M0I | N/A | Random Intercept at Mouse Level | 4575.8 | 4592.1 | -2284.9 | 3 | N/A | N/A | N/A | N/A |
| Structural | M0 | M0I + *test* + *sex* + *sugar* + *exp* + *blockDay* | | 4497.2 | 4562.4 | -2236.6 | 13 | M0I | 10 | 96.58 | $2.2 \times 10^{-16}$ * |
| Core Scientific | M1 | M0 + *line*temp*conc* | | 4129.8 | 4233.0 | -2045.9 | 20 | M0 | 7 | 381.39 | < 0.0001 * |
| *line:exp* interaction | M2a | M1 + *line:exp* | | 4124.2 | 4232.9 | -2042.1 | 21 | M1 | 1 | 7.56 | 0.0060 * |
| *temp:exp* interaction | M2b | M1 + *temp:exp* | | 4131.8 | 4240.5 | -2045.9 | 21 | M1 | 1 | < 0.01 | 0.9919 |
| *conc:exp* interaction | M2c | M1 + *conc:exp* | | 4129.6 | 4238.3 | -2044.8 | 21 | M1 | 1 | 2.17 | 0.1411 |
| All 2-way *exp* interactions | M2d | M1 + *line:exp* + *temp:exp* + *conc:exp* | | 4125.8 | 4245.3 | -2040.9 | 23 | M1<br>M2a<br>M2b<br>M2c | M1: 3<br>M2a: 2<br>M2b: 2<br>M2c: 2 | M1: 10.03<br>M2a: 2.47<br>M2b: 10.03<br>M2c: 7.86 | M1: 0.0183 *<br>M2a: 0.2902<br>M2b: 0.0066 *<br>M2c: 0.0196 * |
| *temp:conc:exp* interactions | M2e | M1 + *temp:conc:exp* | | 4133.4 | 4252.9 | -2044.7 | 23 | M1<br>M2b<br>M2c | M1: 3<br>M2b: 2<br>M2c: 2 | M1: 2.40<br>M2b: 2.40<br>M2c: 0.23 | M1: 0.4936<br>M2b: 0.3012<br>M2c: 0.8896 |
| *line:exp* + *conc:exp* interactions | M2f | M1 + *line:exp* + *conc:exp* | | 4123.8 | 4237.9 | -2040.9 | 22 | M1<br>M2a<br>M2c | M1: 2<br>M2a: 1<br>M2c: 1 | M1: 10.02<br>M2a: 2.47<br>M2c: 7.86 | M1: 0.0067 *<br>M2a: 0.1163<br>M2c: 0.0051 * |
| ***line:conc:exp* interactions** | **M2g** | **M1 + *line:conc:exp*** | | **4120.8** | **4240.3** | **-2038.4** | **23** | **M1<br>M2a<br>M2c** | **M1: 3<br>M2a: 2<br>M2c: 2** | **M1: 15.00<br>M2a: 7.44<br>M2c: 12.83** | **M1: 0.0018 *<br>M2a: 0.0242 *<br>M2c: 0.0016 *** |

| | | | | | | | | | | |
|---|---|---|---|---|---|---|---|---|---|---|
| | | | | | | | **M2f** | **M2f: 1** | **M2f: 4.97** | **M2f: 0.0257 *** |
| *line:sex* (M1) | M3a | M1 + *line:sex* | 4131.2 | 4239.9 | -2045.6 | 21 | M1 | 1 | 0.55 | 0.4581 |
| *line:sex* (M2g) | M3b | M2g + *line:sex* | 4122.2 | 4247.2 | -2038.1 | 24 | M2g | 1 | 0.57 | 0.4496 |
| *test:blockDay* (M1) | M4a | M1 + *test:blockDay* | 4136.7 | 4267.1 | -2044.4 | 30 | M1 | 10 | 3.09 | 0.6868 |
| *test:blockDay* (M2g) | M4b | M2g + *test:blockDay* | 4127.9 | 4274.6 | -2037.0 | 33 | M2g | 10 | 2.89 | 0.7172 |
| *line:blockDay* (M1) | M4c | M1 + *line:blockDay* | 4131.6 | 4240.3 | -2045.8 | 22 | M1 | 2 | 0.20 | 0.6566 |
| *line:blockDay* (M2g) | M4d | M2g + *line:blockDay* | 4122.8 | 4247.8 | -2038.4 | 25 | M2g | 2 | < 0.01 | 0.9999 |

**Table S3** - Model fit parameters and outcomes of likelihood-ratio tests used in model fitting and final model selection for the sugar data. The final analytical model (M2g) is bolded. Total observations = 1692; Total subjects = 47; AIC: Aikake Information Criterion; BIC: Bayesian Information Criterion; DF: Degrees of Freedom; LL: Log-Likelihood; N/A: Not Applicable. * = $p < 0.05$

| Model Term | Reference Level | Predictor Level | Estimate | SE | p-value |
|---|---|---|---|---|---|
| $\beta_0$ | All terms at reference level (*exp* = 0) | Intercept (Baseline Estimate) | 0.791 | 0.219 | 0.0003 * |
| $\beta_{test2}$ | Reference Test (Table 2: Test 1) | Table 2: Test 2 | -0.247 | 0.080 | 0.0020 * |
| $\beta_{test3}$ | | Table 2: Test 3 | 0.096 | 0.073 | 0.1862 |
| $\beta_{test4}$ | | Table 2: Test 4 | 0.133 | 0.073 | 0.0671 |
| $\beta_{test5}$ | | Table 2: Test 5 | 0.012 | 0.073 | 0.8638 |
| $\beta_{test6}$ | | Table 2: Test 6 | -0.007 | 0.073 | 0.9182 |
| $\beta_{line}$ | B6 | KO | 0.072 | 0.225 | 0.7495 |
| $\beta_{sex}$ | Female | Male | 0.196 | 0.196 | 0.3177 |
| $\beta_{sugar}$ | Glucose | Sucrose | 0.049 | 0.196 | 0.8009 |
| $\beta_{temp}$ | 15°C | 30°C | -0.034 | 0.085 | 0.6842 |
| $\beta_{conc}$ | 100 mM | 500 mM | 0.951 | 0.125 | $3.9 \times 10^{-14}$ * |
| $\beta_{blockDay1}$ | $1^{st}$ day of a Test block | $2^{nd}$ day of a Test block | -0.032 | 0.046 | 0.4821 |
| $\beta_{blockDay2}$ | | $3^{rd}$ day of a Test block | -0.073 | 0.046 | 0.1174 |
| $\beta_{exposure}$ | $1^{st}$ Exposure (*exp* = 0) | $2^{nd}$ Exposure (*exp* = 1) | 0.010 | 0.007 | 0.1844 |
| $\beta_{line:temp}$ | B6, 15°C | KO, 30°C | -0.242 | 0.106 | 0.0227 * |
| $\beta_{line:conc}$ | B6, 100 mM | KO, 500 mM | -0.443 | 0.165 | 0.0073 * |
| $\beta_{temp:conc}$ | 15°C, 100 mM | 30°C, 500 mM | -0.035 | 0.120 | 0.7706 |
| $\beta_{line:exposure}$ | B6, Exposure 0 | KO, Exposure 1 | -0.004 | 0.010 | 0.7033 |
| $\beta_{conc:exposure}$ | 100 mM, Exposure 0 | 100 mM, Exposure 1 | 0.028 | 0.010 | 0.0071 * |
| $\beta_{line:temp:conc}$ | B6, 15°C, 100 mM | KO, 30°C, 500 mM | 0.017 | 0.150 | 0.9116 |
| $\beta_{line:conc:exposure}$ | B6, 100 mM, Exposure 0 | KO, 500 mM, Exposure 1 | -0.033 | 0.015 | 0.0257 * |

**Table S4** - Fixed effect coefficient estimates from the final analytical sugar linear mixed model. Note that all values are reported in log+1 scale. Estimates, and their standard errors and p-values, are shown for the reference-coded model terms predicting lick responses. Reference Level indicates the reference group for that term. The Estimate represents the estimated change in log+1 scale licks between a Predictor and its Reference Level (or 0, for Intercept), and the p-value the significance of that effect. Intercept represents the estimated response at all Reference Levels and exposure = 0 (i.e. Day 1). SE: Standard Error. * = $p < 0.05$

| Context // Reference | 0.03 mM @ 15°C | 0.03 mM @ 30°C | 0.3 mM @ 15°C | 0.3 mM @ 30°C |
|---|---|---|---|---|
| 0.03 mM @ 15°C | – | -24.56 [-36.21, -12.91] | -23.08 [-34.77, -11.38] | -35.98 [-47.79, -24.16] |
| 0.03 mM @ 30°C | -6.13 [-17.78, 5.52] | – | -17.26 [-29.07, -5.44] | -27.97 [-39.62, -16.31] |
| 0.3 mM @ 15°C | -8.16 [-19.85, 3.53] | -20.77 [-32.60, -8.96] | – | -30.38 [-42.03, -18.73] |
| 0.3 mM @ 30°C | -2.63 [-14.44, 9.19] | -13.05 [-24.71, -1.40] | -11.95 [-23.60, -0.30] | – |

**Table S5** - Estimated marginal mean water-adjusted licks (WAL) [95% confidence interval] for a given reference solution (columns) when paired alongside each of its three context solutions (rows). Note, a lower value indicates greater aversion.

| Context // Reference | 100 mM @ 15°C | 100 mM @ 30°C | 500 mM @ 15°C | 500 mM @ 30°C |
|---|---|---|---|---|
| 100 mM @ 15°C | – | 2.81 [2.08, 3.76] | 10.24 [8.06, 12.75] | 7.45 [5.88, 9.42] |
| 100 mM @ 30°C | 3.45 [2.66, 4.49] | – | 9.34 [7.45, 11.75] | 8.73 [6.89, 11.06] |
| 500 mM @ 15°C | 3.90 [2.99, 4.98] | 2.86 [2.14, 3.76] | – | 5.65 [4.43, 7.14] |
| 500 mM @ 30°C | 3.42 [2.56, 4.41] | 3.35 [2.50, 4.36] | 6.98 [5.48, 8.89] | – |

**Table S6**- Estimated marginal mean licks [95% confidence interval] for a given reference solution (columns) when paired alongside each of the other three context solutions (rows).
